# Bilateral Intracortical Inhibition during Unilateral Motor Preparation and Sequence Learning

**DOI:** 10.1101/2023.10.19.563212

**Authors:** R. Hamel, B. M. Waltzing, M.R. Hinder, C. McAllister, N. Jenkinson, J.M Galea

**Author notes:** Correspondence should be addressed to Dr Raphael Hamel.

## Abstract

Motor sequence learning gradually quickens reaction time, suggesting that sequence learning alters motor preparation processes. Interestingly, evidence has shown that preparing sequence movements decreases short intracortical inhibition (SICI) in the contralateral motor cortex (M1), but also that sequence learning alters motor preparation processes in both the contralateral and ipsilateral M1s. Therefore, one possibility is that sequence learning alters the SICI decreases occurring during motor preparation in bilateral M1s. To examine this, two novel hypotheses were tested: unilateral sequence preparation would decrease SICI in bilateral M1s, and sequence learning would alter such bilateral SICI responses. Paired-pulse transcranial magnetic stimulation was delivered over the contralateral and ipsilateral M1s to assess SICI in an index finger muscle during the preparation of sequences initiated by either the right index or little finger. In the absence of sequence learning, SICI decreased in both the contralateral and ipsilateral M1s during the preparation of sequences initiated by the right index finger, suggesting that SICI decreases in bilateral M1s during unilateral motor preparation. As sequence learning progressed, SICI decreased in the contralateral M1 whilst it increased in the ipsilateral M1. Moreover, these bilateral SICI responses were observed at the onset of motor preparation, suggesting that sequence learning altered baseline SICI levels rather than the SICI decreases occurring during motor preparation *per se*. Altogether, these results suggest that SICI responses in bilateral M1s reflect two motor processes: an acute decrease of inhibition during motor preparation, and a cooperative but bidirectional shift of baseline inhibition levels as sequence learning progresses.

## INTRODUCTION

Motor sequence learning is a paradigm in which a repeating sequence of movements is performed with increasing speed and accuracy^1–3^. Interestingly, work has shown that sequence learning reduces reaction time (RT) by altering the processes involved in preparing the sequences^4,5^. For instance, Ariani and Diedrichsen (2019)^4^ showed that constraining the time permitted to prepare individual sequences slows motor sequence learning, suggesting that motor preparation processes contribute to sequence learning. One key question is what neural process (or processes) of motor preparation is (are) altered as sequence learning progresses?

To address this question, one must consider the known processes of motor preparation, neural substrates of sequence learning, and whether sequence learning alters motor preparation in these substrates. First, brain stimulation evidence suggests that one robust process of motor preparation consists of short intracortical inhibition (SICI) decreases in the contralateral motor cortex (M1)^6–8^, which is thought to reflect decreases in intracortical type A gamma-aminobutyric acid (GABA_A_)-mediated inhibition^9^. For instance, using paired-pulse transcranial magnetic stimulation (ppTMS), Hamel et al. (2023)^8^ showed that SICI decreases in the left M1 as participants prepare to execute sequences with their right hand. Given that sequence learning alters motor preparation processes^4,5^, one possibility is that sequence learning alters such SICI decreases during motor preparation^8^. Moreover, other ppTMS studies complement this evidence by showing that resting SICI levels decrease in the contralateral M1 when learning to perform sequential pinch^10^ and fast thumb abduction movements^11^, which further hints at the possibility that SICI levels are altered as sequence learning progresses. Second, TMS studies have shown that both the contralateral^12,13^ and ipsilateral^14–17^ M1s causally contribute to motor sequence learning. For instance, Kobayashi et al. (2009)^15^ showed that inhibitory repetitive TMS (rTMS) delivered before sequence learning disrupts the RT quickening when applied over the contralateral M1, but enhances the quickening of RT when applied over the ipsilateral M1, suggesting that processes occurring in bilateral M1s underpin sequence learning. Third, additional neuroimaging^18,19^ and TMS^20^ work importantly extends the above evidence by suggesting that sequence learning *specifically* alters motor preparation processes in bilateral M1s. For instance, Hamano et al. (2021)^18^ recorded functional magnetic resonance imaging data to show that sequence learning increases activity in the contralateral M1 during motor preparation. Moreover, Cohen et al. (2009)^20^ showed that delivering TMS pulses over the ipsilateral M1 during motor preparation impairs the memory consolidation of sequence learning, suggesting that motor preparation processes are not selectively altered in the contralateral M1. Altogether, this evidence suggests that SICI decreases in the contralateral M1 is a neural process of sequence preparation, but also that sequence learning alters preparation processes in both the contralateral and ipsilateral M1s. Consequently, one tantalising possibility is that sequence learning alters the SICI decreases occurring during motor preparation in bilateral M1s, and not selectively in the contralateral M1.

To examine this possibility, this study tested two hypotheses. The first hypothesis was that, in the absence of sequence learning, SICI in M1 would decrease bilaterally during the preparation of unilateral sequence movements. This hypothesis is based on single-pulse TMS studies showing that unilateral motor preparation increases corticospinal excitability (CSE) in both the contralateral (see ref ^21^ for a review) and ipsilateral M1s^22^, suggesting that bilateral changes in M1s’ excitability underpin motor preparation (see ref ^23^ for further support). This first hypothesis was expected to confirm that SICI decreases in bilateral M1s during sequence preparation, therefore providing a trackable motor preparation process as sequence learning progresses. Based on the results of this first hypothesis and on evidence suggesting that sequence learning alters motor preparation processes in bilateral M1s^18–20^, the second hypothesis was that sequence learning would alter SICI in bilateral M1s during motor preparation.

## Methods

### Participants

Four groups of 20 medication-free, self-reported neurologically healthy and right-handed young adults participated in this study. Overall, the 80 participants were 20.4 ± 0.5 years old (51 females and 29 males). Hereafter, all descriptive statistics represent the mean ± 95% confidence intervals (CIs). Participants were screened for TMS contraindications^24^ and provided their informed written consent (which was approved by the local institutional board; project # ERN_17-1541AP6). All procedures were in accordance with the Declaration of Helsinki. Participants received course credits in exchange for their enrolment in this study.

### Overview of the procedures

Two experiments were performed using ppTMS to assess SICI as participants prepared to execute unilateral right-handed 4-element finger-press sequences. Each experiment contained a Left M1 and a Right M1 group. In the Left M1 groups, TMS was applied over the left (contralateral) M1 whilst MEPs were collected from a right (task-relevant) index finger muscle. Conversely, in the Right M1 groups, TMS was applied over the right (ipsilateral) M1 whilst MEPs were collected from a left (task-irrelevant) index finger muscle.

The first experiment (Experiment #1; Figure 1) tested the first hypothesis, which was that *in the absence* of sequence learning, SICI in M1 would decrease bilaterally during unilateral motor preparation. Specifically, SICI was assessed in the contralateral (Left M1 group; n = 20) and ipsilateral M1s (Right M1 group; n = 20) during the preparation of pseudorandomised sequences, which was to prevent sequence learning. The second experiment (Experiment #2; Figure 2) tested the second hypothesis, which was that *the presence* of sequence learning would alter the SICI decreases occurring during motor preparation in bilateral M1s. Specifically, SICI was also assessed in the contralateral (Left M1 group; n = 20) and ipsilateral M1 (Right M1 group; n = 20) during unilateral motor preparation, but this time *in the presence* of sequence learning.

**Figure 1.**
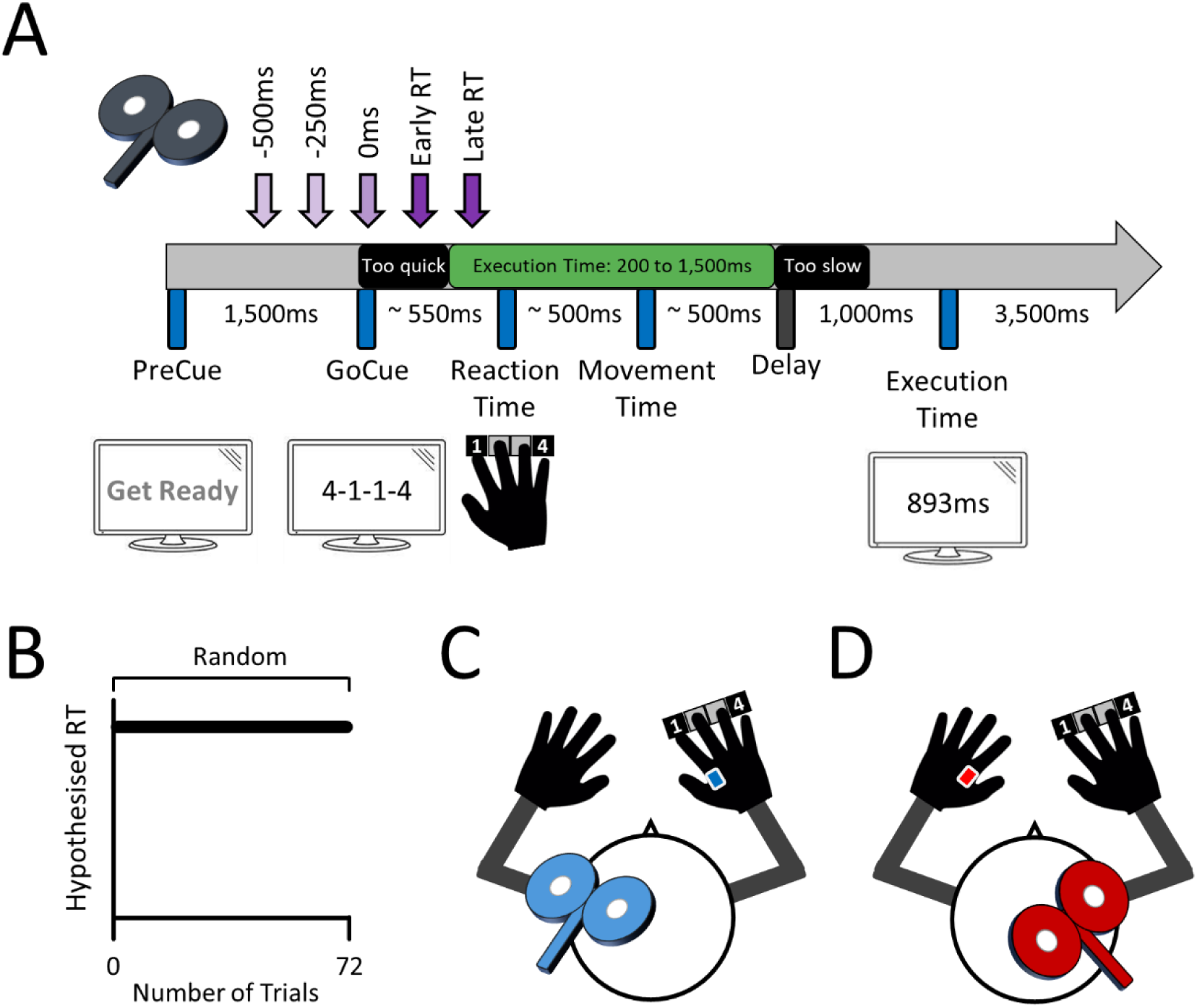
Experiment one – TMS delivery over the left and right M1 during motor preparation in the absence of sequence learning. **(A)** *Chronology of a typical trial –* In response to a visual GoCue, participants used their right hand to execute 4-element finger-press sequences as quickly and accurately as possible. TMS pulses were delivered either before the GoCue (−500ms, −250ms), at GoCue (0ms), or after the GoCue (during the RT period; +250ms, +500ms). The two TMS time points after the GoCue were (re)pooled according to the latency at which pulses were delivered relative to the RT (Early RT: ≥ 1% but ≤ 50% of the RT; Late RT: > 50% but ≤ 100% of the RT). **(B)** *Random blocks –* The finger-press sequences were pseudorandomised to prevent sequence learning, and to isolate the effects of motor preparation on CSE and SICI. Participants performed a total of 4 Random blocks (72 trials per block; 1 sequence per trial). **(C)** *Left M1 group* – TMS was applied over the left M1 and the data were recorded from the right (task-relevant) first dorsal interosseous (FDI) muscle. **(D)** *Right M1 group* – TMS was applied over the right M1 and the data were recorded from the left (task-irrelevant) FDI muscle.

**Figure 2.**
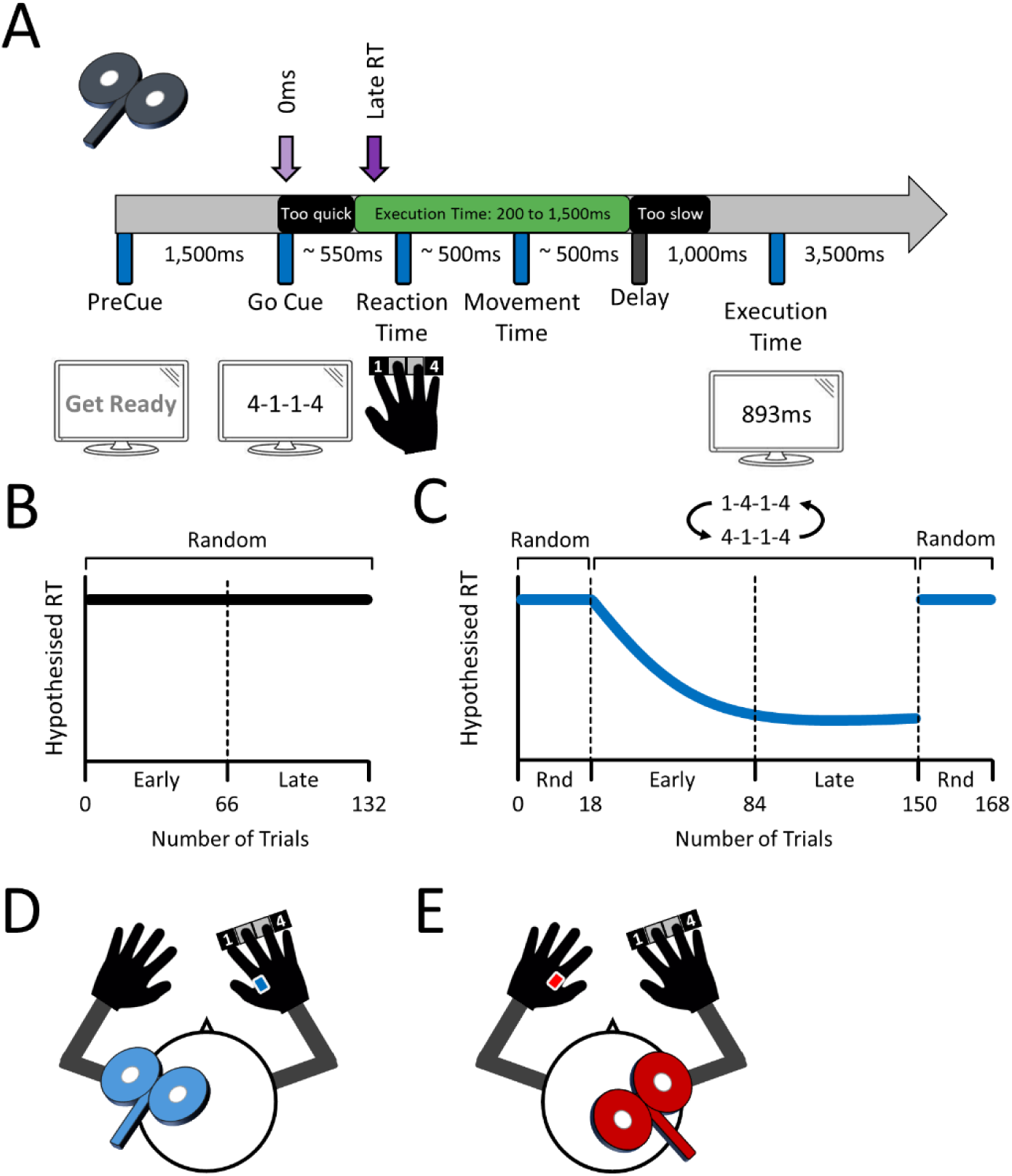
Second experiment – TMS delivery over the left and right M1 during motor preparation as sequence learning was induced. **(A)** *Chronology of a typical trial* – Based on the findings of the first experiment, TMS pulses were selectively delivered at GoCue and Late RT during motor preparation. This was to increase the number of TMS trials per time point. Unlike the first experiment, the Late RT time point was iteratively calculated as 75% of the RT median based on a sliding window, which was to account for the gradual RT quickening as sequence learning progresses. **(B)** *Random blocks* – To prevent sequence learning (control condition), participants executed pseudorandomised sequences. **(C)** *Learning blocks* – To induce sequence learning, participants executed sets of 2 repeating sequences. For both Random and Learning blocks, the first and second half of trials were also grouped and respectively labelled Early and Late phases. Each participant performed a total of 2 Random and 2 Learning blocks in a fully counterbalanced order. **(D)** *Left M1 group –* TMS was applied over the contralateral M1 and the data were recorded from the right (task-relevant) FDI muscle. **(E)** *Right M1 group –* TMS was applied over the ipsilateral M1 and the data were recorded from the left (task-irrelevant) FDI muscle.

### Finger-press sequences, apparatus and trial chronology

In both experiments, the participants executed sequences comprising index and little finger presses of their right hand on a USB-wired keyboard (600 Microsoft ®). The “D” and “J” keyboard keys were labelled as digits “1” and “4”, and corresponded to index and little finger presses, respectively. Designing sequences using these two digits only resulted in 6 possible finger-press sequences: “1-1-4-4”, “1-4-1-4”, “1-4-4-1”, “4-4-1-1”, “4-1-4-1”, and “4-1-1-4”.

All visual stimuli were displayed on a 24-inch Iiyama Prolite (B2409HDS) computer monitor (1920 x 1080 pixels; 60Hz vertical refresh rate). The behavioural experiments were run through custom-built scripts using MATLAB (R2021b; MathWorks ®) and the PsychToolBox-3 interface. The use of the PsychToolBox-3 function “KbQueueCheck” resulted in sub-millisecond sampling (> 1,000Hz) for both the finger-press timing and the latency of the TMS pulses. A USB-wired Arduino Nano board with a Deek Robot Terminal Adapter was controlled through MATLAB to externally trigger the TMS devices. Based on 800 behavioural trials and bootstrapped estimations (100,000 samples), the Arduino hardware had an average latency of ∼20 ± 6ms between issuing the MATLAB command and the actual trigger delivery. This was offset by sending the trigger 20ms earlier than the predefined time points. The 3ms inter-pulse interval was defined in, and controlled by, Signal (Cambridge Electronic Design, v6.05), ensuring that the Arduino hardware latencies did not alter the SICI pulses’ inter-pulse interval. Finally, the SICI pulses were also triggered a further 3ms earlier, to offset the inter-pulse interval. This was to ensure that test pulses would be delivered at a similar latency on single and paired-pulse trials to evaluate CSE and SICI.

The trial chronology was identical across both experiments (Figures 1A and 2A). A trial was initiated with the presentation of a PreCue (“Get Ready”) for 1,500ms. The GoCue then displayed the finger-press sequence to be executed. A total of 1,750ms was allowed to prepare and execute the sequence (“Allowed Execution Time” in Figures 1A and 2A). Once four keys were pressed or the 1,750ms limit was reached, whichever came first, the screen went black for a delay of 1,000ms. The execution time of the ongoing trial was displayed for 1,500ms, after which the trial ended with a black screen. Each trial lasted ∼8,000ms with a fixed inter-trial interval of 2,000ms.

### Design of each experiment

Experiment one examined the effects of motor preparation on SICI *in the absence* of sequence learning (Motor Preparation; Figure 1B). To avoid sequence learning, participants performed the six finger-press sequences in a pseudorandomised order over 4 blocks of 72 trials. Each of these blocks, labelled as ‘random’ contained six repeats of a 12-trial cycle, where each of the six possible finger sequences was presented twice. There was an even split of sequences initiated with the index and little fingers, and the same sequence never repeated on adjacent trials.

Each 12-trial cycle contained 10 TMS and 2 NoTMS trials. Over the 10 TMS trials, each single-(CSE) and paired-pulse (SICI) TMS variable was assessed at each of the 5 Time Points (−500ms, −250ms, GoCue, +250ms, and +500ms). The NoTMS trials were to evaluate behavioural performance in the absence of the disruptive effects of TMS^25,26^. Halfway through each Random block, participants were provided with 2-minute breaks to relieve fatigue. A total of 288 trials (240 TMS and 48 NoTMS trials) were collected for both the Left and Right M1 groups, which necessitated approximately 40 min. From start to finish, the first experiment lasted approximately 65 minutes.

Experiment two examined the effects of motor preparation on SICI *in the presence* of sequence learning (Sequence Learning; Figure 2). Participants performed a total of 2 Learning and 2 Random blocks in a fully counterbalanced order. As in experiment one, the Random blocks (Figure 2B) contained pseudorandomised sequences to prevent sequence learning, therefore acting as a control for the Learning blocks. Each Random block contained 132 trials split into three 44-trial cycles. Within each 44-trial cycle, each TMS variable (CSE, SICI) was assessed at each of the 2 Time Points (GoCue, Late RT) a total of 6 times (36 TMS trials). Each 44-trial cycle also contained 8 NoTMS trials. Participants were provided with 2-minute breaks to relieve fatigue at the end of each 44-trial cycle. In each Random block, there was an even split of sequences initiated by the index and little fingers, and the same sequence never repeated on adjacent trials. When combining the 2 Random blocks, a total of 264 trials (216 TMS and 48 NoTMS trials) were collected for both the Left M1 and Right M1 groups, which necessitated approximately 35 minutes.

Each Learning block (Figure 2C) started and ended with short blocks of pseudorandomised sequences. Each of these consisted of 18 NoTMS trials, and the first short block was used to individualise the Late RT time point (see below) as sequence learning was induced. To induce sequence learning, sets of 2 repeating sequences were executed for a total of 132 trials (e.g., first Learning block: “1-4-1-4 → 4-1-1-4”; second Learning block “4-1-4-1” → “1-4-4-1”). During those 132 trials, three 44-trial cycles were designed, in which the sets of two repeating sequences were executed a total of 22 times. As for the Random blocks, for each 44-trial cycle, each TMS variable (SICI, CSE) was assessed at each of the 2 time points (GoCue, Late RT) a total of 6 times (36 TMS trials). Each cycle also contained 8 NoTMS trials. Participants were provided with 2-minute breaks to relieve fatigue at the end of each 44-trial cycle. In each Learning block, there was an even split of sequences initiated by the index and little fingers. When combining the two Learning blocks, a total of 336 trials (216 TMS and 120 NoTMS trials) were collected for the Left M1 and Right M1 groups, which necessitated approximately 50 min. From start to finish, the second experiment lasted approximately 110 minutes.

### Justification of the TMS Time Points

In the first experiment (Motor Preparation), TMS pulses were delivered at 5 Time Points (Figure 1A) to investigate excitability changes before the GoCue (during the foreperiod), at GoCue (onset of motor preparation), and after GoCue (during motor preparation)^6,7,27^. Specifically, TMS was either delivered at −500ms or −250ms before the GoCue, at the GoCue (0ms), or +250ms or +500ms after the GoCue. After they were collected, the data from the +250ms and +500ms time points were pooled as a function of the latency at which TMS pulses were delivered relative to the actual reaction time (RT) *in that trial*. Specifically, pulses delivered ≥ 1% but ≤ 50% of the RT period were pooled and identified as Early RT. Pulses delivered > 50% but ≤ 100% of the RT period were pooled and identified as Late RT. Hereafter, the +250ms and +500ms time points of the first experiment are referred to as Early RT and Late RT, respectively.

In the second experiment (Sequence Learning), excitability changes were only assessed at GoCue (0ms) and Late RT time points (Figure 2A). Reducing the number of time points maximised the number of TMS trials at GoCue and Late RT to ensure robust estimates of excitability. Unlike in the first experiment, the latency of the Late RT time point was iteratively adjusted to correspond to 75% of the RT period, which was to control for the progressive quickening of RT as sequence learning progressed. Specifically, in Learning blocks, the first 18 NoTMS trials were used as a baseline to calculate 75% of the median RT, determining the latency of the first trial on which TMS pulses would be delivered at Late RT. Then, as sequence learning was ongoing, the value corresponding to 75% of the median RT was iteratively calculated using a sliding window spanning the previous 18 trials, which was used to adjust the latency of the Late RT time point. This procedure was applied throughout each Learning block, except for the last 18 trials where no TMS pulses were delivered. In Random blocks, an identical procedure to control the latency of the Late RT time point was used, except for the first 18 trials where TMS pulses were delivered. Namely, in the first 18 trials, TMS pulses delivered at Late RT assumed a +500ms latency. After the 18^th^ trial, this latency was iteratively adjusted by using the same procedure as above for the whole duration of a Random block, including the last 18 trials where TMS pulses were delivered. For both the Random and Learning blocks, the latency of the pulses delivered at GoCue was not iteratively adjusted, as they coincided with the start of motor preparation.

A posteriori descriptive analyses confirmed that the sliding median window procedure was successful at individualising the latency of the Late RT time point to ∼75% of the RT period. For the Left M1 group, Late RT pulses occurred at a latency of 76.0 ± 0.4% and 76.1 ± 0.4% of the RT period for the Learning and Random blocks, respectively. For the Right M1 group, Late RT pulses occurred at RT latencies of 76.3 ± 0.4 and 75.3 ± 0.4% for the Learning and Random blocks, respectively.

### EMG system and transcranial magnetic stimulation

Electromyography (EMG) activity was recorded from the right and left FDI (Figures 1C-D and 2D-E) using a 2-channel Delsys Bagnoli system (Delsys, Natick, USA). Importantly, the FDI is a co-agonist of the index finger flexions required for performing button presses^28^, indicating its contribution to executing the present sequences. It was reasoned that if FDI activity would increase when the index or the little finger initiated the sequences, this would allow determining if the observed excitability changes were specific to the finger that initiated the sequences. Hereafter, the term “finger-specific” refers to excitability changes observed selectively when the index or little finger initiated the sequences. Oppositely, the term “finger-unspecific” refers to excitability changes observed when both the index and little fingers initiated the sequences.

The EMG data from the FDI muscles were acquired with the Signal software (v6.05) and a Micro 1401 analogue-to-digital converter (both Cambridge Electronic Design, Cambridge, UK). EMG signals were sampled at 10,000Hz for epochs of 500ms (200ms pre-trigger time). The EMG data were band-pass filtered between 20 and 450Hz, with a notch filter applied at 50Hz. The reference EMG electrode was positioned on the proximal olecranon process of the ulnar bone of the same limb. The EMG data were analysed using an automated custom-built MatLab script.

Neuronavigated TMS pulses were delivered over M1 through a single figure-of-eight 70mm Alpha Flat Coil (taped) connected to a paired-pulse BiStim^2^ stimulator (MagStim, Whitland, UK)^29^. The coil was positioned at a 45° angle in a posterior-anterior axis over the FDI motor hotspot. The motor hotspot was defined as the cortical area in M1 where motor-evoked potentials (MEPs) of maximal amplitude could be reliably elicited with suprathreshold pulses in the FDI. The resting motor threshold (RMT) was defined as the percentage (%) of maximum stimulator output to induce 5 out of 10 MEPs of at least 50µV of peak-to-peak amplitude^30^. The test stimulus (TS) intensity was set to induce MEPs of ± 1mV at rest. For each participant, the FDI motor hotspot, the RMT, and the TS intensity were assessed at the start of the TMS session. Brainsight (Rogue Research; Montreal, Canada) ensured reliable coil positioning during all experimental procedures^29^.

To assess CSE, single test pulses of TMS at TS intensity were delivered. To assess SICI, the conditioning pulse was set at the intensity corresponding to 70% of the RMT^31^, the inter-pulse interval was set at 3ms^32^, and the test pulse was set at TS intensity. To confirm that these TMS parameters could successfully induce SICI in both the left and right M1s, 30 CSE and 30 SICI pulses were delivered at rest before participants began the experiments. As described below, SICI was evaluated and normalised to CSE and generalised mixed models were used to analyse the data. In the first experiment, SICI pulses induced inhibition in both the Left (39.1 ± 6.7%; χ^2^ = 40.71; *p* < 0.0001) and Right M1 groups (32.1 ± 4.2%; χ^2^ = 65.39; *p* < 0.0001) as compared to CSE pulses (100%). Similarly, in the second experiment, SICI pulses induced inhibition in both the Left (39.4 ± 6.2%; χ^2^ = 40.71; *p* < 0.0001) and Right M1 groups (44.7 ± 8.5%; χ^2^ = 42.10; *p* < 0.0001) as compared to CSE pulses (100%). For both experiments, this confirms that SICI was reliably induced at rest in both the contralateral and ipsilateral M1s.

### Number of TMS trials per condition

In both experiments, a total of 24 TMS trials were delivered for each TMS variable (CSE, SICI) and Time Point, except for the second experiment (Sequence Learning) where a total of 30 TMS trials for each TMS variable were delivered at Late RT. Recording a total of 24 TMS trials per time point provides a robust estimation of SICI and CSE^33,34^. Recording a total of 30 TMS trials at Late RT in the second experiment was to compensate for the added variability in RTs as sequence learning was ongoing, which was expected to result in more trial rejection at that time point. Overall, for both experiments, approximately 21 valid TMS trials were included on average for each TMS variable and time point, suggesting robust estimations of CSE and SICI^33,34^. See Supplementary Table 1 for a complete report of the final number of valid TMS trials included in the analyses.

### Dependent variables

In both experiments, RT was defined as the time difference in ms between GoCue onset and the first finger press. Movement time (MT) was defined as the time difference in ms between the first and last finger press. Accuracy was defined as the correct sequence execution within the allowed 1,750 msec execution time and was measured as a binary variable (1 = success, 0 = failure). To evaluate CSE, MEP peak-to-peak amplitudes induced by single test pulses were calculated as (non-normalized) values in mV for each time point. To evaluate SICI, the MEP peak-to-peak amplitudes induced by the SICI test pulses were first calculated for each time point. Then, these individual SICI trials were normalised as a percentage (%) of their corresponding average CSE.

### Trial rejection and data preparation for analyses

In both experiments, RTs below 150ms were rejected to avoid including trials with premature responses. To prevent the contamination of muscle pre-activation on MEP amplitudes, trials where EMG activity with a root mean square exceeding 100µV in the 50ms before the TMS pulse was delivered were rejected (similar to Smith et al., 2019)^35^. Importantly, TMS pulses that were delivered with a latency > 100% of the RT were rejected, since TMS pulses would then be delivered during execution (outside of motor preparation). In the second experiment only, TMS pulses delivered at Late RT with a latency of < 50% of the RT were also rejected. This was to ensure that only trials delivered in the second half (> 50% but < 100%) of the RT period were included at Late RT.

To evaluate the effects of TMS pulses on behaviour during the first experiment (Motor Preparation), the RT and MT data were normalised to the NoTMS trials. Specifically, RT and MT data of individual trials were divided by the average RT and MT from the NoTMS trials, therefore resulting in a trial-per-trial percent (%) change. This was performed separately for each participant. The binomial variable accuracy could not be normalised to the average of NoTMS trials because divisions by zero are impossible. Specifically, the mean accuracy from CSE and SICI trials was directly compared to that of NoTMS trials.

To evaluate changes in behavioural and TMS data during the second experiment (Sequence Learning), data from each Learning and Random block were split into an early (first 66 trials) and late phase (last 66 trials; Figure 2B-C). This trial separation was to ensure enough valid TMS trials (at least 21) for each phase, therefore providing robust estimates of CSE and SICI as sequence learning progressed^33,34^. In the second experiment, all the RT, MT, Accuracy, CSE and SICI data were pooled into an Early and Late phase.

### Statistical analyses

In both experiments, generalised mixed models^36,37^ with a gamma distribution to account for the positive continuous skewness were conducted to analyse RT, MT, CSE, and SICI data^38^. To analyse the binomial Accuracy data, generalised mixed models with a binomial distribution were conducted. For each model, the maximally complex random effect structure (random intercepts for Participants and random slopes for all of the fixed effects and interactions) that minimised the Akaike Information Criterion was chosen to analyse the results^39^.

In the first experiment (Motor Preparation), the fixed effects to analyse the behavioural data were 5 Time Points (−500ms, −250ms, 0ms, Early RT, Late RT) * 3 TMS Pulse Types (NoTMS, CSE, SICI). The fixed effects to analyse the TMS data were 5 Time Points (−500ms, −250ms, 0ms, Early RT, Late RT) * 2 Initiating Fingers (Index, Little), where CSE and SICI data were analysed separately. In the second experiment (Sequence Learning), the fixed effects to analyse the behavioural data were 2 Blocks (Random, Learning) * 2 Phases (Early, Late) * 3 TMS Pulse Types (NoTMS, CSE, SICI). The fixed effects to analyse the TMS data were 2 Blocks (Random, Learning) * 2 Phases (Early, Late) * 2 Time Points (GoCue, Late RT) * 2 Initiating Fingers (Index, Little), where CSE and SICI data were analysed separately.

For all analyses, *p-values* below 0.05 were determined as statistically significant. The Benjamini-Hochberg (1995) correction was used to correct *p* values for multiple comparisons^40^. As mentioned above, the mean ± 95% CIs were used to report statistics throughout. All analyses were conducted in JAMOVI (version 2.3.16)^41^.

### Data availability statement

All data are freely available at the following URL: https://osf.io/tc3dx/

## Results – First Experiment (Motor Preparation)

### Motor preparation: Finger-specific CSE increases selective to the contralateral M1

For the Left M1 group (i.e., task-relevant FDI), the CSE data (Figure 3A) revealed a Time Points * Initiating Fingers interaction (χ^2^ = 22.58, *p* < 0.001), which was decomposed by conducting simple effects of Time Points (−500ms, −250ms, GoCue, Early RT, Late RT) at each level of Initiating Fingers (Index, Little). When the right Index finger initiated the sequences, CSE increased at Late RT as compared to all the other time points (all *p* < 0.001). When the right Little finger initiated the sequences, CSE *decreased* at Late RT as compared to all the other time points (all *p* < 0.001). This shows that as movement onset approached, CSE in projections to the right FDI *increased* when initiating the sequences with the right index finger, whereas CSE *decreased* when initiating the sequences with the little finger on the same right hand.

**Figure 3.**
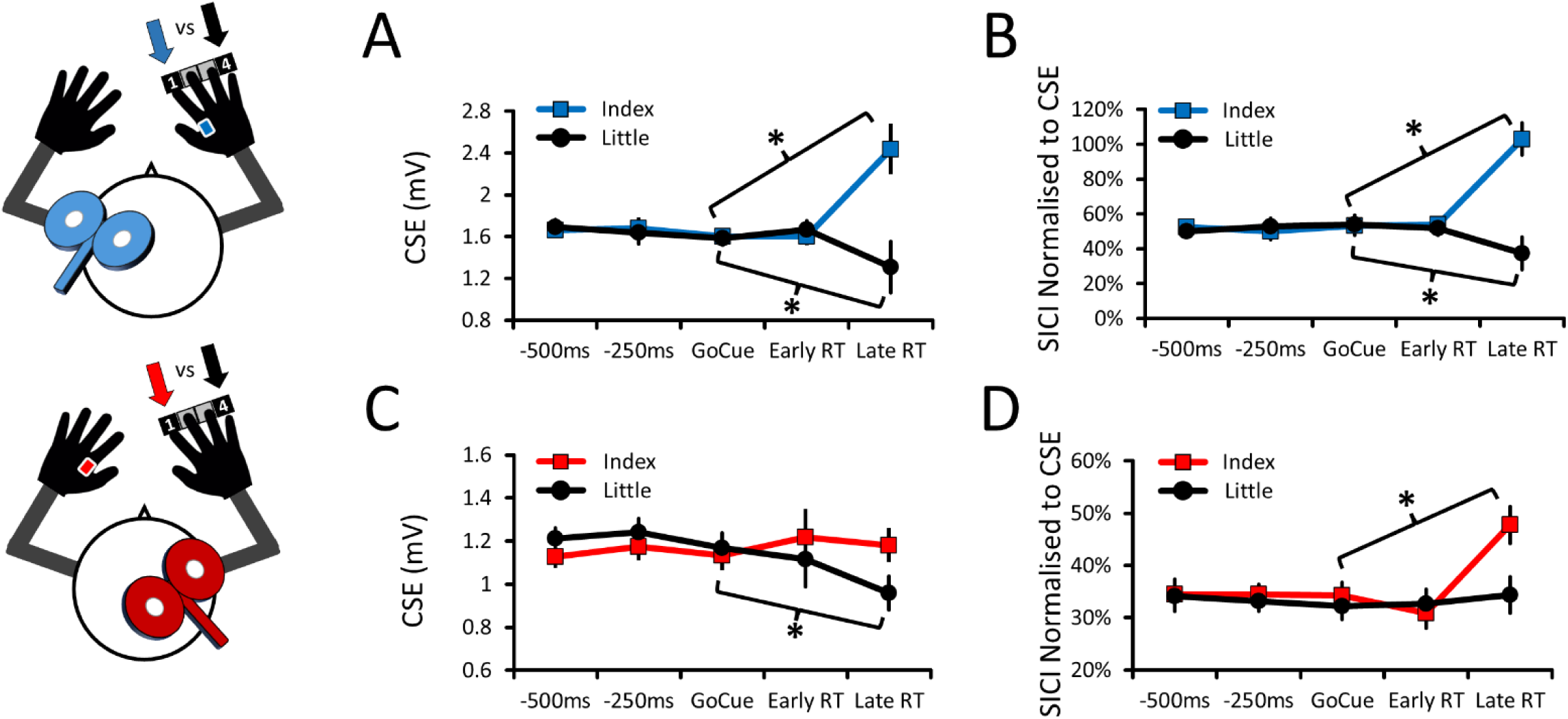
CSE and SICI data when TMS was delivered to the left and right M1 during preparation. For all panels, the data are pooled according to the finger (Index vs Little) of the right hand that initiated the sequences. **(A)** *Left M1 group* – CSE increased in the right FDI when the right Index initiated the sequences but decreased when the right Little finger did. **(B)** *Left M1 group* – SICI decreased in the right FDI when the right Index initiated the sequences but increased when the right Little finger did. **(C)** *Right M1 group* – CSE did not change in the left FDI when the right Index initiated the sequences but decreased when the right Little finger did. **(D)** *Right M1 group* – SICI decreased in the left FDI when the right Index initiated the sequences but did not change when the right Little finger did. For all panels, the mean and within-subject 95% CIs are shown. The asterisks (*) denote significant differences between the GoCue and Late RT time points (*p* < 0.05). To enhance clarity and maintain consistency with the time points of the second experiment, only the differences between GoCue and Late RT are identified above.

For the Right M1 group (i.e., task-irrelevant FDI), the CSE data (Figure 3C) also revealed a Time Points * Initiating Fingers interaction (χ^2^ = 22.29, *p* < 0.001), which was decomposed as above. When the Index finger initiated the sequences, CSE did not change at Late RT as compared to all the other time points (all *p* > 0.627). However, when the Little finger initiated the sequences, CSE *decreased* at Late RT as compared to all the other time points (all *p* < 0.002). This shows that as movement onset approached, CSE in projections to the left FDI did not change when initiating the sequences with the right (homologous) index finger, whereas CSE *decreased* when initiating the sequences with the right little finger.

Altogether, these results suggest that movement-onset-related CSE increases in the FDI are specific to the contralateral M1 during the preparation of sequences initiated by the right index finger. Moreover, the results suggest that movement-onset-related CSE *decreases* in the FDI occur in bilateral M1s during the preparation of sequences initiated by the right little finger.

### Motor preparation: Finger-specific SICI decreases in bilateral M1s

For the Left M1 group (i.e., task-relevant FDI), the SICI data (Figure 3B) revealed a Time Points * Initiating Fingers interaction (χ^2^ = 135.95, *p* < 0.001), which was decomposed as above. When the Index finger initiated the sequences, SICI decreased at Late RT as compared to all the other time points (all *p* < 0.001). When the Little finger initiated the sequences, SICI *increased* at Late RT as compared to all the other time points (all *p* < 0.002). This shows that as movement onset approached, SICI in the cortical representation of the right FDI muscle decreased when initiating the sequences with the right index finger, whereas SICI *increased* when initiating the sequences with the little finger of the same right hand.

For the Right M1 group (i.e., task-irrelevant FDI), the SICI data (Figure 3D) revealed a Time Points * Initiating Fingers interaction (χ^2^ = 13.40, *p* = 0.010), which was also decomposed as above. When the Index finger initiated the sequences, SICI decreased at Late RT as compared to all the other time points (all *p* < 0.008). When the Little finger initiated the sequences, SICI did not change at Late RT as compared to all the other time points (all *p* > 0.847). This shows that as movement onset approached, SICI in the cortical representation of the left FDI muscle decreased when initiating the sequences with the right (homologous) index finger, whereas SICI did not change when initiating the sequences with the right little finger.

Altogether, these results suggest that movement-onset-related SICI decreases in the FDI occur in bilateral M1s during the preparation of sequences initiated by the right index finger. Furthermore, this also suggests that information about finger-specificity is shared between the contralateral and ipsilateral M1s. As apparent in Figure 3B-3D, it should be noted that the magnitude of the SICI decreases at Late RT in the Right M1 (∼50%) was approximately half of those in the Left M1 (∼100%). Finally, the results also suggest that movement-onset-related SICI *increases* in the right FDI are selective to the contralateral M1 when initiating the sequences with the right little finger.

### Motor preparation: Effects of TMS over the Left and Right M1 on sequence performance

For full details on how TMS pulses over the contralateral and ipsilateral M1 during preparation altered the RT, MT, and Accuracy of the right hand as compared to NoTMS trials, see the Supplementary Results and Supplementary Figure 1. Briefly, in both the Left M1 and Right M1 groups, TMS pulses quickened RTs to a similar extent when delivered during the foreperiod (−500ms, −250ms) and at the onset of motor preparation (0ms), and also similarly slowed RTs when delivered during preparation (Early RT). In contrast, TMS pulses delivered late during motor preparation (Late RT) slowed MTs and decreased Accuracy, but only in the Left M1 group. Overall, this suggests that the processes of motor preparation are similarly altered by TMS pulses over bilateral M1s, whereas the processes of motor execution are selectively disrupted by TMS pulses over the contralateral M1.

## Results – Second Experiment (Sequence Learning)

### Sequence learning: RTs quickened in Learning but not in Random blocks

For the Left M1 group, the RT data (Figure 4A) selectively revealed a Blocks * Phases (χ^2^ = 12.39; *p* < 0.001), which was decomposed by conducting simple effects of Phases (Early vs Late) separately at each level of Blocks (Random, Learning). See Supplementary Table 2 for a complete report of all results. In the Learning blocks, RTs quickened from the Early to Late phase (χ^2^ = 15.14; *p* < 0.001), but not in Random ones (χ^2^ = 0.58; *p* = 0.445). This confirms the presence of sequence learning in the Left M1 group. This also suggests that delivering TMS pulses over the contralateral M1s did not alter sequence learning.

**Figure 4.**
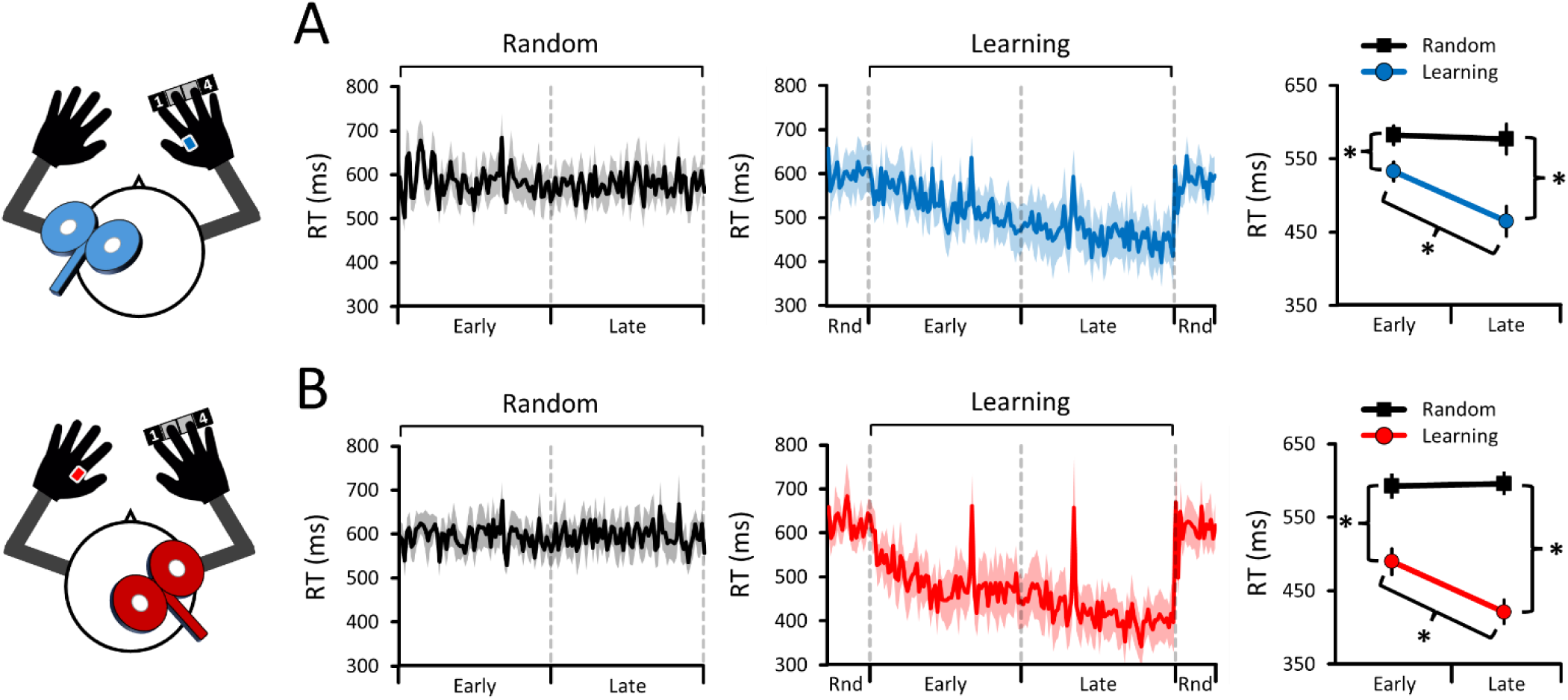
RT data as sequence learning with the right hand progressed. **(A)** *Left M1 group* – RTs quickened in the Learning blocks (repeating sequences) but not in the Random ones (pseudorandomised sequences). **(B)** *Right M1 group* – RTs quickened in the Learning blocks but not in the Random ones. Overall, this confirms the presence of sequence learning in both the Left M1 and Right M1 groups. Moreover, as compared to NoTMS trials, delivering TMS pulses to the contralateral M1 did not prevent sequence learning, but delivering TMS pulses to the ipsilateral M1 quickened RTs (not shown in Figure 4). For all panels, the mean and within-subject 95% CIs are shown (either as shaded or whisker error bars). The asterisks (*) denote significant differences (*p* < 0.05). “Rnd” means Random (pseudorandomised sequences).

For the Right M1 group, the RT data (Figure 4B) revealed a Blocks * Phases (χ^2^ = 56.57; *p* < 0.001) and Blocks * TMS Pulse Types interactions (χ^2^ = 6.17, *p* = 0.046). See Supplementary Table 3 for a complete report of all results. As for the Left M1 group, RTs quickened from the Early to Late phase in the Learning blocks (χ^2^ = 52.50, *p* < 0.001), but not in the Random ones (χ^2^ = 0.17, *p* = 0.677). Moreover, the Blocks * TMS Pulse Types interaction revealed that, in both Random (χ^2^ =13.40, *p* = 0.001) and Learning blocks (χ^2^ = 51.63, *p* < 0.001), both CSE (both *p* < 0.009) and SICI pulses (both *p* < 0.001) *quickened* RTs as compared to NoTMS. This confirms the presence of sequence learning in the Right M1 group, and suggests that TMS pulses over the ipsilateral M1 during motor preparation can enhance the RT quickening during sequence learning (similar to ^15^).

### Sequence learning: MTs quickened in Learning blocks, but Accuracy did not change

For full details of the MT and accuracy results, see the Supplementary Results, Supplementary Tables 4 to 7, and Supplementary Figure 2. Briefly, for both the Left and Right M1 groups, MTs quickened in the Learning blocks, suggesting that sequence learning did not solely quicken RTs. However, in both groups, Accuracy did not change as sequence learning progressed, suggesting that the RT and MT improvements were not accompanied by changes in accuracy. As in the first experiment, TMS pulses over the contralateral M1, but not the ipsilateral one, slowed MTs and decreased accuracy as compared to NoTMS trials.

### Motor preparation: Finger-specific CSE increases selective to the contralateral M1

For the Left M1 group (i.e., task-relevant FDI), the CSE data revealed a Block * Time Points * Initiating Fingers interaction (χ^2^ = 26.08; *p* < 0.001; Figure 5A), which was decomposed by conducting simple effects of Time Points (GoCue vs Late RT) at each level of Blocks (Random, Learning) and Initiating Fingers (Index, Little). See Supplementary Table 8 for a complete report of all results. When the right Index finger initiated the sequences, CSE increased from GoCue to Late RT in both the Random (χ^2^ = 49.71, *p* < 0.001) and Learning blocks (χ^2^ = 29.61, *p* < 0.001). When the right Little finger initiated the sequences, the results revealed that CSE *decreased* from GoCue to Late RT in the Random blocks (χ^2^ = 24.27, *p* < 0.001), but not in Learning ones (χ^2^ < 0.01, *p* = 0.993). These results replicate the finger-specific CSE responses observed in the Left M1 group of the first experiment (Random blocks).

**Figure 5.**
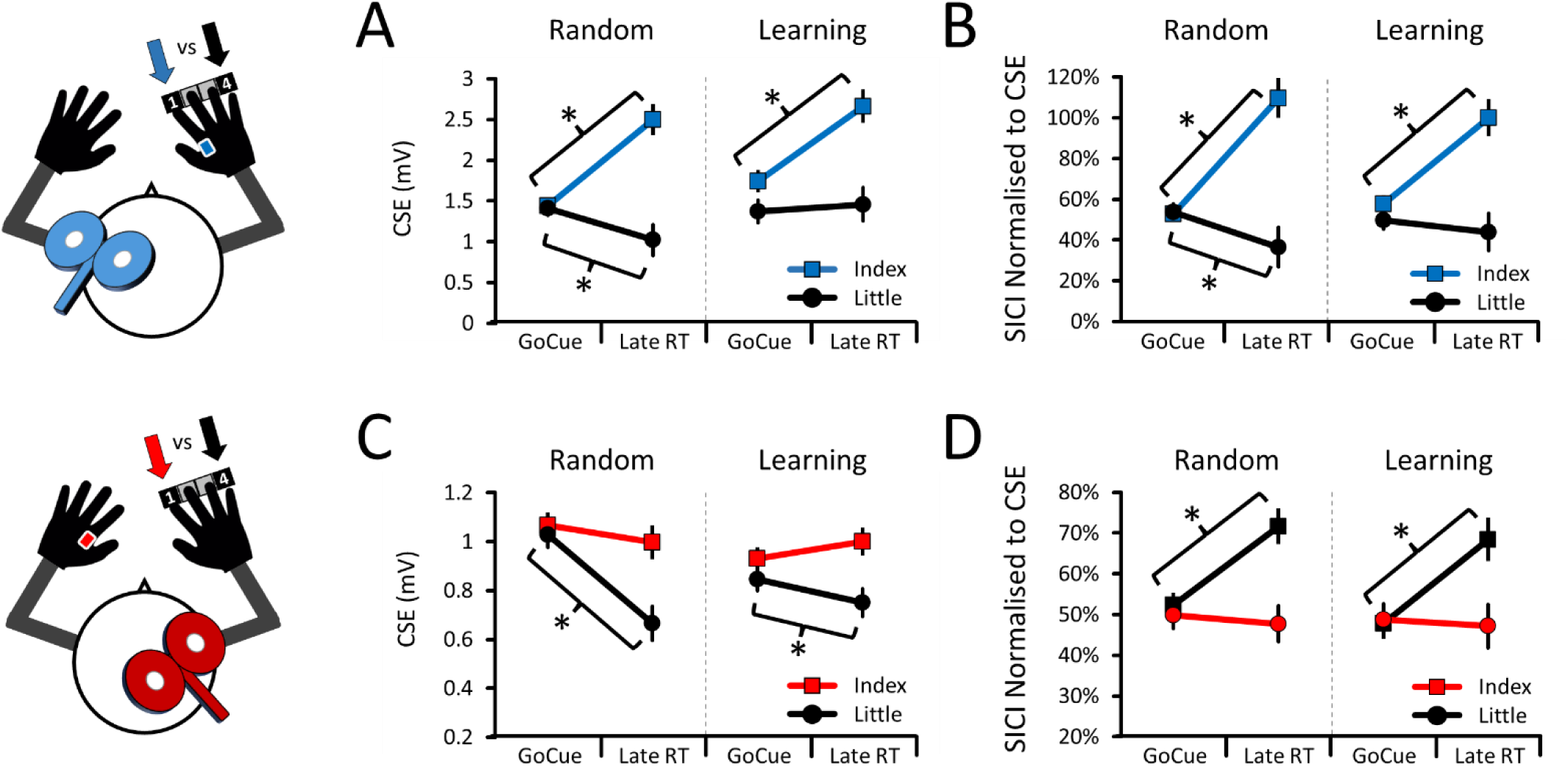
Finger-specific CSE and SICI data during motor preparation. For all panels, the data are pooled according to the finger (Index vs Little) of the right hand that initiated the sequences. **(A)** *Left M1 group* – CSE increased in the right FDI when the right Index initiated the sequences in both the Random and Learning blocks. Initiating the sequences with the right Little finger decreased CSE in the Random blocks only. **(B)** *Left M1 group* – SICI decreased in the right FDI when the right Index initiated the sequences in both the Random and Learning blocks. Initiating the sequences with the right Little finger increased SICI in the Random blocks only. **(C)** *Right M1 group* – CSE did not change in the left FDI when the right Index but decreased when the right Little finger initiated the sequences in both the Random and Learning blocks. **(D)** *Right M1 group* – SICI decreased in the left FDI when the right Index, but did not increase when the right Little finger, initiated the sequences in both the Random and Learning blocks. For all panels, the mean and within-subject 95% CIs are shown. The asterisks (*) denote significant differences between the GoCue and Late RT time points (*p* < 0.05).

For the Right M1 group (i.e., task-irrelevant FDI), the CSE data also revealed a Blocks * Time Points * Initiating Fingers interaction (χ^2^ = 3.82; *p* = 0.051; Figure 5C), which was decomposed as above. See Supplementary Table 9 for a complete report of all results. When the right Index finger initiated the sequences, CSE did not increase from GoCue to Late RT in both the Random (χ^2^ = 0.39; *p* = 0.531) and Learning blocks (χ^2^ = 0.62; *p* = 0.431). When the right Little finger initiated the sequences, CSE *decreased* from GoCue to Late RT in the Random (χ^2^ = 24.13; *p* < 0.001) and also tended to decrease in the Learning blocks (χ^2^ = 4.45; *p* = 0.070). As for the Left M1 group, these results also replicate the finger-specific CSE responses observed in the Right M1 group of the first experiment (Random blocks).

### Motor preparation: Finger-specific SICI decreases in bilateral M1s

For the Left M1 group (i.e., task-relevant FDI), the SICI data revealed a Blocks * Time Points * Initiating Fingers interaction (χ^2^ = 6.37, *p* = 0.012; Figure 5B), which was decomposed as above. See Supplementary Table 10 for a complete report of all results. When the right Index finger initiated the sequences, SICI decreased from GoCue to Late RT in both the Random (χ^2^ = 40.16, *p* < 0.001) and Learning blocks (χ^2^ = 55.07, *p* < 0.001). When the right Little finger initiated the sequences, SICI *increased* from GoCue to Late RT in the Random blocks (χ^2^ = 9.72, *p* = 0.002), but not in Learning ones (χ^2^ = 0.31, *p* = 0.576). Overall, these results replicate the finger-specific SICI responses observed in the Left M1 group of the first experiment (Random blocks).

For the Right M1 group (i.e., task-irrelevant FDI), the SICI data revealed a Time Points * Initiating Fingers interaction (χ^2^ = 16.29; *p* < 0.001; Figure 5D), which was decomposed by conducting simple effects of Time Points (GoCue vs Late RT) at each level of Initiating Fingers (Index, Little). See Supplementary Table 11 for a complete report of all results. When the Index finger initiated the sequences, SICI decreased from GoCue to Late RT (χ^2^ = 0.31, *p* = 0.576). When the Little finger initiated the sequences, SICI did not change from GoCue to Late RT (χ^2^ = 1.18; *p* = 0.278). Note that data in Figure 5D were further separated by Random and Learning blocks to maintain visual consistency with the other panels. Overall, these results replicate the finger-specific SICI responses observed in the Right M1 group of the first experiment (Random blocks). Finally, as apparent in Figure 5B-5D, it should be noted that the magnitude of the SICI decreases at Late RT in the Right M1 (∼70%) was smaller than those observed in the Left M1 (∼105%).

### Sequence learning: CSE was not altered in the contralateral or ipsilateral M1s

For the Left M1 group (i.e., task-relevant FDI), the CSE data revealed no Blocks * Time Points * Phases interaction (χ^2^ = 1.92; *p* = 0.166; Figure 6A), suggesting that CSE in projections to the right FDI was not altered as sequence learning progressed in the right hand. Similarly, for the Right M1 group (i.e., task-irrelevant FDI), the CSE data also revealed no Blocks * Time Points * Phases (χ^2^ = 2.65; *p* = 0.103; Figure 6C), suggesting that CSE in projections to the left FDI was also not altered as sequence learning progressed in the right hand. See Supplementary Tables 8 and 9 for a complete report of all results. Overall, this shows that sequence learning with the right hand did not alter CSE in both the contralateral and ipsilateral M1s (i.e., in both task-relevant and task-irrelevant FDI).

**Figure 6.**
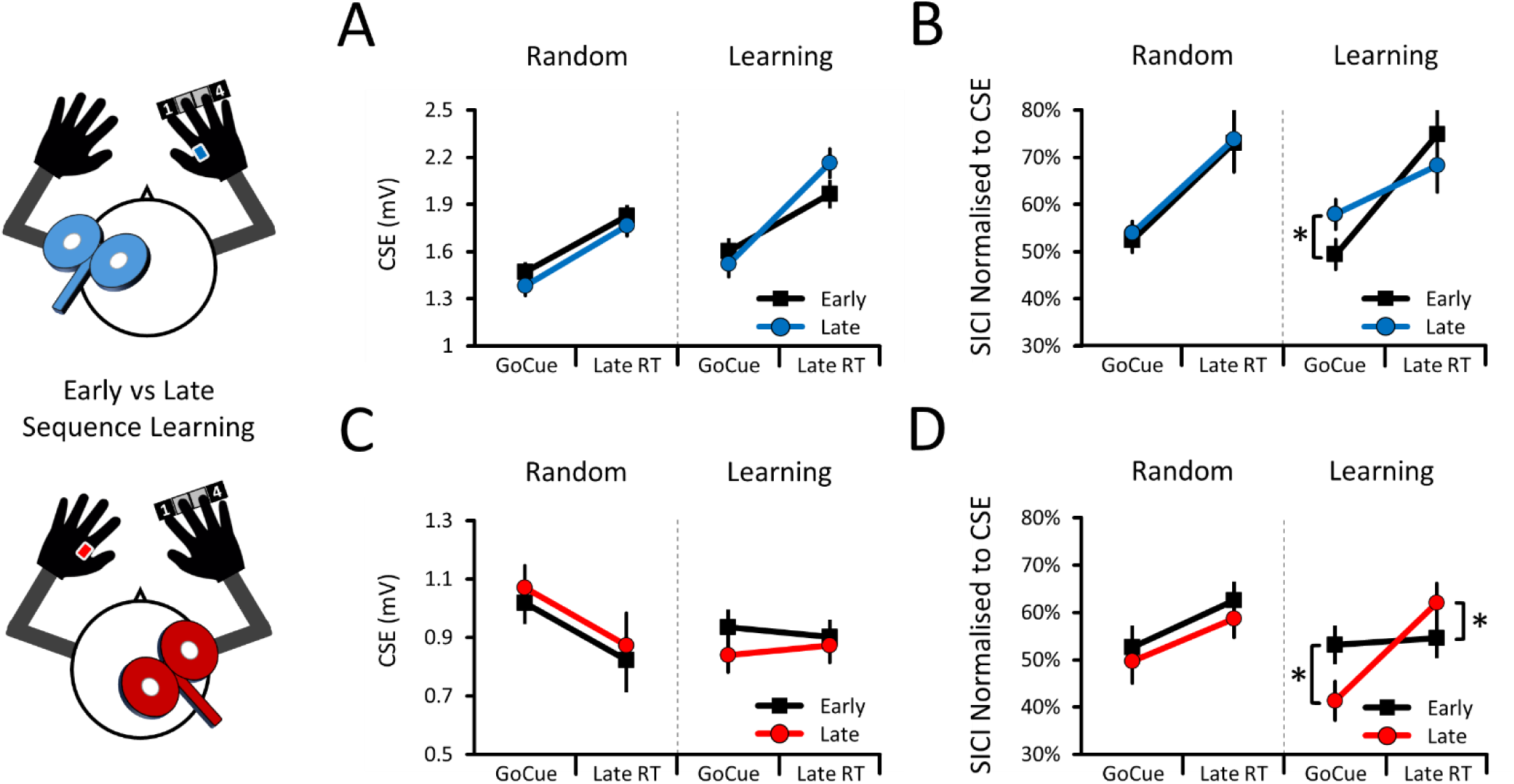
Phase-specific CSE and SICI data during preparation as sequence learning with the right hand progressed. For all panels, the data are pooled according to the Phases (Early vs Late) of sequence learning. **(A)** *Left M1 group* – CSE did not change in the right FDI from the Early to Late phase for both the Random and Learning blocks. **(B)** *Left M1 group* – SICI decreased in the right FDI at GoCue from the Early to Late phase in the Learning blocks only. Although not significant, SICI tended to increase from the Early to Late phase at Late RT (*p* = 0.125). **(C)** *Right M1 group* – CSE did not change in the left FDI from the Early to Late phase for both Random and Learning blocks. **(D)** *Right M1 group* – SICI increased in the left FDI at GoCue from the Early to Late phase in Learning blocks only. Also, SICI decreased at Late RT from the Early to Late phase in Learning blocks only. For all panels, the mean and within-subject 95% CIs are shown. Asterisks (*) denote significant differences between the Early and Late phases (*p* < 0.05).

### Sequence learning: SICI decreased in the contralateral M1 whilst increasing in the ipsilateral M1 at GoCue

For the Left M1 group (i.e., task-relevant FDI), the SICI data revealed a Blocks * Time Points * Phases interaction (χ^2^ = 3.94, *p* = 0.047; Figure 6B), which was decomposed by conducting simple effects of Phases (Early vs Late) at each level of Blocks (Random, Learning) and Time Points (GoCue, Late RT). The Random blocks revealed that SICI did not change from the Early to Late phase at both GoCue (χ^2^ = 0.30, *p* = 0.583) and Late RT (χ^2^ = 0.04, *p* = 0.840). However, the Learning blocks revealed that SICI *decreased* from the Early to Late phase when measured at GoCue (χ^2^ = 7.56, *p* = 0.001), but not at Late RT (χ^2^ = 2.35, *p* = 0.125). This shows that sequence learning with the right hand decreased SICI in the cortical representation of the right FDI, but at the onset of motor preparation only. The results also show that SICI did not change in the Random blocks, suggesting learning-specificity. Finally, the results also show that SICI decreased both when the index and little fingers initiated the sequences, suggesting that sequence learning decreased SICI in a finger-unspecific manner.

For the Right M1 group (i.e., task-irrelevant FDI), the SICI data also revealed a Blocks * Time Points * Phases interaction (χ^2^ = 8.89; *p* = 0.003; Figure 6D), which was decomposed as above. The Random blocks revealed that SICI did not change from the Early to Late phase at both GoCue (χ^2^ = 0.05, *p* = 0.825) and Late RT (χ^2^ = 3.11, *p* = 0.156). However, the Learning blocks revealed that SICI *increased* from the Early to Late phase at GoCue (χ^2^ = 5.30, *p* = 0.021), but that SICI *decreased* at Late RT (χ^2^ = 4.60, *p* = 0.032). This shows that sequence learning with the right-hand altered SICI in the cortical representation of the left FDI. Unlike for the Left M1 group, this shows that SICI increased at GoCue but decreased at Late RT, suggesting that the SICI responses in the ipsilateral M1 were present both at the onset of, and *during,* motor preparation. As for the Left M1 group, the results show that these SICI responses were learning-specific, and finger-unspecific.

Altogether, these results suggest that sequence learning with the right-hand decreased SICI in the contralateral M1 whilst it increased SICI in the ipsilateral M1. Moreover, the finding that these SICI responses were observed in both M1s at GoCue, but not at Late RT, suggests that bilateral M1s predominantly alter their tonic baseline SICI levels, rather than the SICI decreases observed during motor preparation *per se* (Figure 3B-D). However, the presence of bidirectional SICI responses during motor preparation cannot be ruled out, as SICI decreased at Late RT in the ipsilateral M1, but also tended to increase in the contralateral M1 (*p* = 0.125).

## Discussion

The two experiments reported here were designed to test two hypotheses: that SICI in M1 would decrease bilaterally during the unilateral preparation of sequence movements, and that sequence learning would alter these bilateral SICI responses. To this end, ppTMS was delivered over the left and right M1 to assess SICI in the right and left FDI muscles, respectively, as participants prepared to initiate sequences with the right index and little fingers. As movement onset approached, SICI decreased bilaterally but selectively when the right index finger initiated the sequences (Figure 3, also shown in summary Figure 7). These results were replicated in the second experiment (Figure 5) and suggest that SICI decreases in bilateral M1s reflect the preparation of unilateral finger-specific motor commands. Conversely, as sequence learning with the right hand progressed, SICI decreased in the contralateral M1 whilst it increased in the ipsilateral M1 (Figure 6, also shown in summary Figure 8). These bilateral and bidirectional SICI responses were learning-specific and observed at the start of the preparation period (i.e., GoCue), suggesting that sequence learning predominantly altered baseline SICI levels rather than the SICI decreases occurring during motor preparation *per se*. Overall, one possibility is that bilateral changes in SICI levels reflect at least two motor processes; the acute SICI decreases as movement onset approaches could support the preparation of motor commands, whereas bidirectional baseline shifts of tonic SICI levels could underpin motor sequence learning.

**Figure 7.**
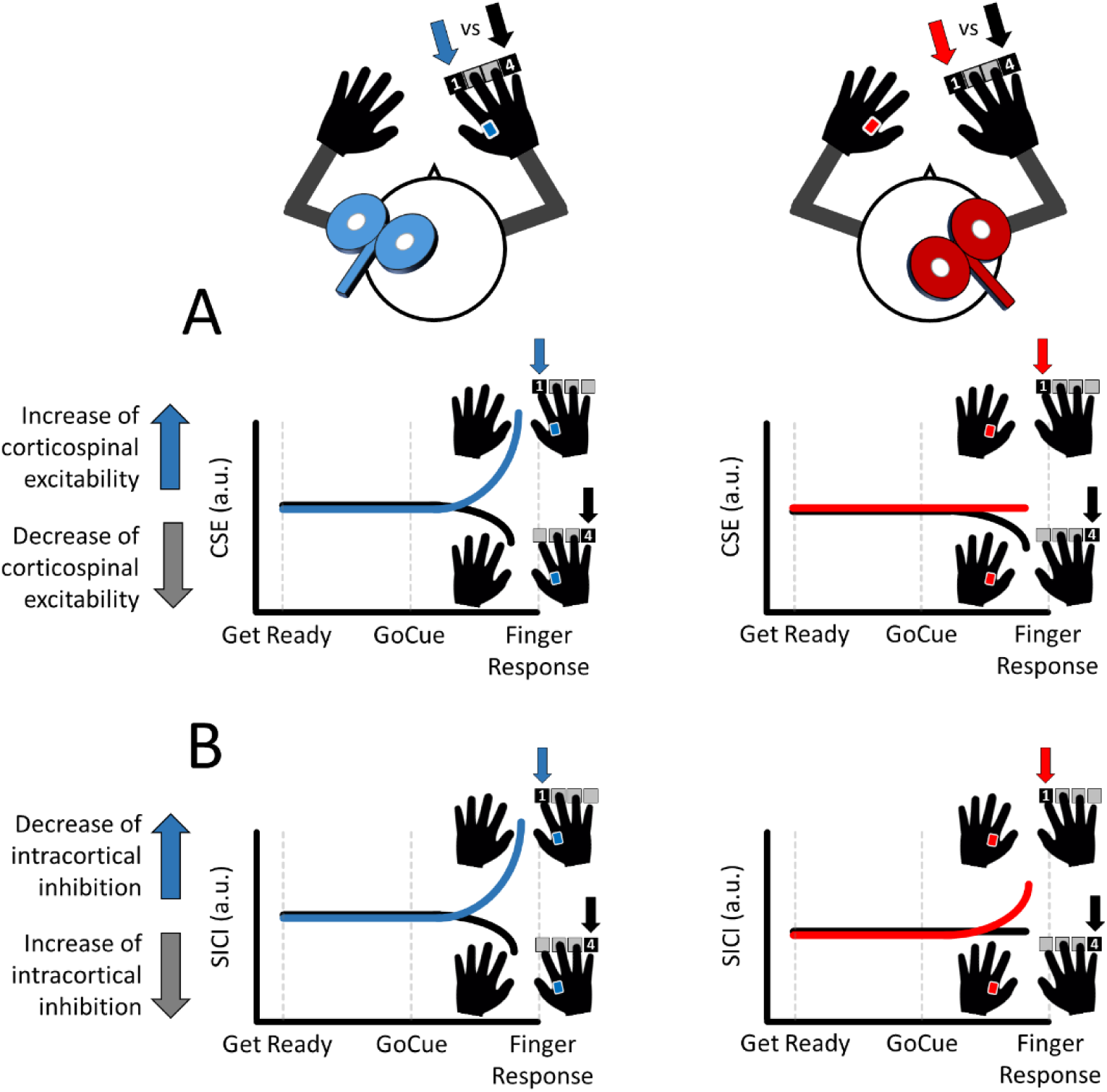
Summary of the key CSE and SICI responses during unilateral motor preparation. **(A)** *CSE results* – Initiating the sequences with the right Index finger increased CSE selectively in the left (contralateral) M1. Initiating the sequences with the right Little finger decreased CSE bilaterally. **(B)** *SICI results* – Initiating the sequences with the right Index finger decreased SICI bilaterally. Initiating the sequences with the right Little finger increased SICI selectively in the contralateral M1.

**Figure 8.**
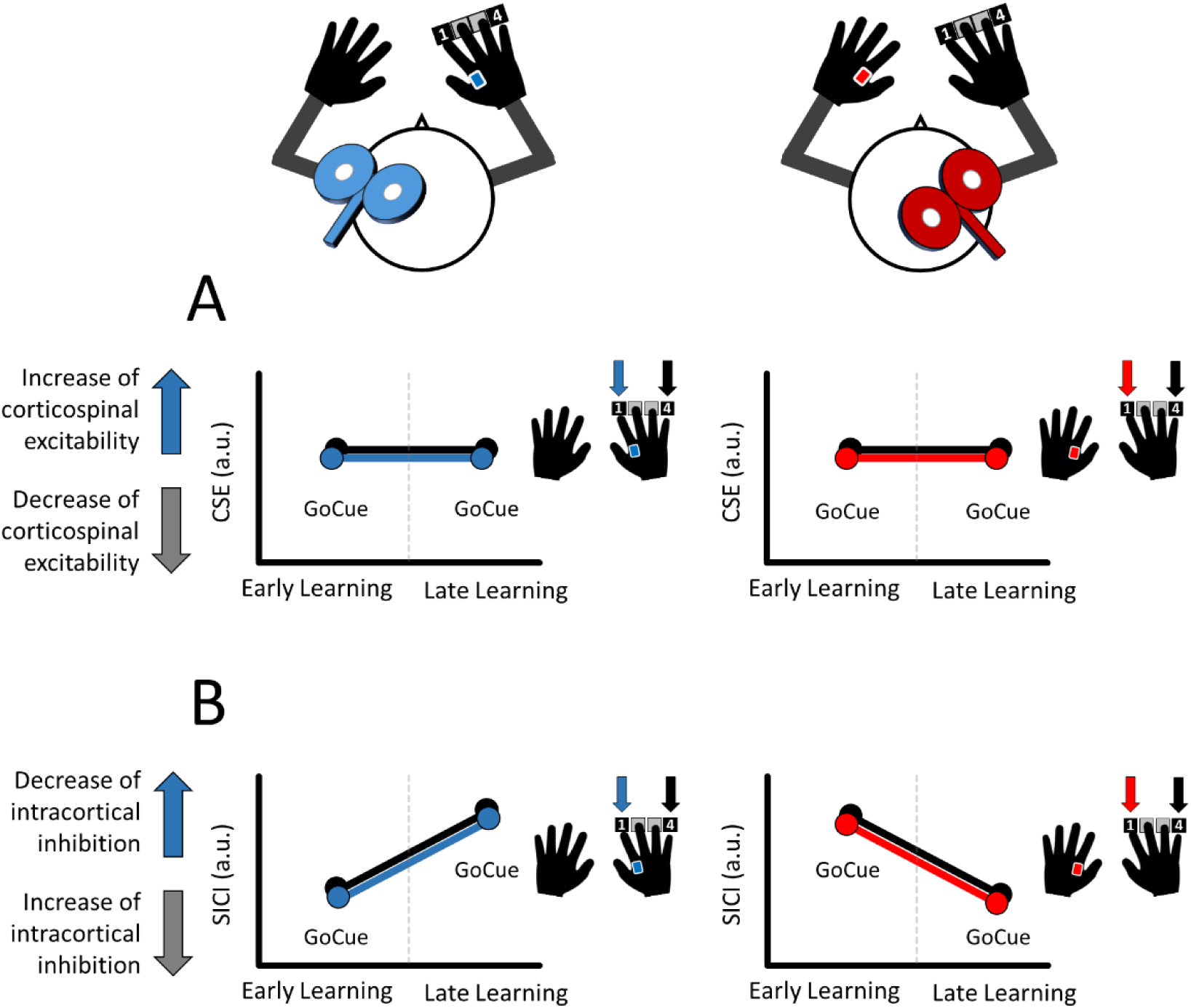
Summary of the key CSE and SICI responses as sequence learning progressed. **(A)** *CSE results* – CSE did neither change in the contralateral nor ipsilateral M1 as sequence learning progressed. **(B)** *SICI results* – As learning progressed, SICI decreased in the contralateral M1 whereas it increased in the ipsilateral one, suggesting that bilateral M1s contribute to sequence learning. This pattern was predominantly observable at GoCue, suggesting that sequence learning selectively altered baseline levels of SICI.

### Bilateral decreases of SICI during unilateral motor preparation

One novel finding is that SICI decreased in bilateral FDI muscles despite sequences being selectively initiated by the right index finger (Figures 3B-D and 5B-D, also shown in summary Figure 7B), suggesting that the preparation of unilateral finger-specific motor commands generates SICI decreases in bilateral M1s. Moreover, the SICI decreases measured before movement onset were of greater amplitude in the contralateral M1 (∼100%) than in the ipsilateral M1 (∼50 to 70%), suggesting a greater contribution from the contralateral M1 to motor preparation (see Chen et al. (1997)^42^ for similar results). Nonetheless, the finding that motor preparation recruits bilateral M1s aligns with previous TMS studies showing contralateral^21^ and ipsilateral^22^ CSE increases as movement onset approaches, and also extends them by suggesting that these previously reported CSE increases originate – at least partly – from SICI decreases in bilateral M1s.

The bilaterality of the SICI decreases, despite movements being unilateral, generates interesting implications. Namely, SICI is considered to reflect local type A gamma-aminobutyric (GABA_A_)-mediated inhibition in M1^9^. As such, one possibility is that preparing unilateral motor commands involves a synchronised cross-hemispheric decrease of local GABA_A_-mediated inhibition in M1. One open question is whether the ipsilateral M1 reflects a copy of the motor commands encoded in the contralateral M1, or if bilateral M1s separately encode distinct motor command parameters^23,42^. Nonetheless, another hint at a cross-hemispheric synchronisation comes from the result that bilateral M1s decreased SICI specifically when the right index finger initiated the sequences, suggesting that some parameters of a unilateral motor command are shared between the contralateral and ipsilateral M1s. Anatomically, such a joint bilateral decrease in SICI could be enabled by homotopic interhemispheric connections between bilateral M1s^43,44^, long-range projection GABAergic interneurons^45^, or could originate from motor areas upstream of M1^46,47^. Although the exact process by which SICI jointly decreases in bilateral M1s during unilateral motor preparation remains unclear, the second experiment replicated this finding in different groups of participants (Figure 5B-D). Thus, one interpretation is that the present bilateral SICI decreases reflect a robust neural process of motor preparation.

An alternative interpretation is that bilateral SICI decreases reflect disinhibition mechanisms required for motor execution^48,49^. In support, previous work showed CSE increases^23,50^ and functional activity increases in bilateral M1s^51–53^ during motor execution, suggesting that execution recruits bilateral M1s^23^. Anatomically, bilateral M1s could contribute to unilateral motor execution through contralateral and ipsilateral corticospinal projections^23^. However, here, CSE increased selectively in the contralateral M1 when the index finger initiated the sequences (Figure 3A, also shown in summary Figure 7A), suggesting that ipsilateral corticospinal projections were not recruited (but opposing previous work^22^). Moreover, as SICI is believed to reflect activity in local intracortical inhibitory circuits^9^, the present bilateral SICI decreases are unlikely to reflect the recruitment of bilateral corticospinal projections. Another hint that preparation could be bilateral but execution could be contralateral comes from the present behavioural results (Supplementary Results). Namely, TMS over both the contralateral and ipsilateral M1s quickened RT to a similar extent when pulses were delivered during the foreperiod and at the onset of motor preparation, and similarly slowed RT when pulses were delivered early during motor preparation. The similarity of these behavioural responses suggests that bilateral M1s similarly contribute to motor preparation processes, although other explanations such as intersensory facilitation cannot be ruled out^54,55^. Nonetheless, TMS pulses delivered late during preparation over the contralateral – but not the ipsilateral – M1 slowed MT and decreased accuracy, suggesting that motor execution processes are contralateralised. Based on this evidence, one possibility is that the present SICI responses – because of their bilaterality – primarily reflect motor preparation processes, as execution could be expected to depend upon contralateralised processes. Future studies using motor imagery^56^ should be able to confirm this interpretation.

It should be noted that the present results do not fully replicate our previous work^8^. Namely, our previous work showed that SICI decreased when both the right index and little fingers initiated the sequences, suggesting finger-unspecificity. In contrast, the present results show that SICI decreased selectively when the right index finger initiated the sequences, rather suggesting finger-specificity. However, our previous work used monetary rewards to incentivise motor performance, which also caused SICI to decrease in a finger-unspecific manner. Therefore, one possibility is that delivering monetary rewards caused finger-specific SICI responses to become finger-unspecific. Nonetheless, in both these studies, the SICI responses were selectively recorded from the FDI muscle, making it unclear how the claims of finger-specificity would extend beyond the index finger.

### Contralateral decrease – then ipsilateral increase – of baseline SICI levels during sequence learning

A key finding is that SICI decreased in the contralateral M1 whilst SICI increased in the ipsilateral M1 as sequence learning progressed from the early to late phase (Figure 6B-D, also shown in summary Figure 8B). Moreover, these SICI responses were absent when sequence learning was prevented, suggesting learning-specificity. One possibility is that bilateral M1s cooperatively contribute to sequence learning by bidirectionally decreasing SICI at different phases of learning.

This finding aligns with mounting evidence suggesting that bilateral M1s cooperate, rather than compete, to support unilateral motor functions^23,50^. One implication of the finding that sequence learning altered bilateral SICI – but not CSE (Figure 6A-C, also shown in summary Figure 8A) – is that such cooperation originates from intracortical rather than corticospinal circuits. Moreover, the bidirectional pattern of SICI responses in bilateral M1s suggests that sequence learning initially induces functional changes in the ipsilateral M1, which then shift to the contralateral M1 as learning progresses. In support, the contralateral M1 has been shown to causally contribute to late, but not early, phases of various motor learning paradigms^57–59^. However, this support is only partial as it remains unknown if the ipsilateral M1 causally contributes to early or late (or both) phases of motor learning^14–17,20^. Nonetheless, the finding that SICI decreased in the contralateral M1 is largely consistent with neuroimaging studies showing that GABA concentrations decrease in the left M1 as sequence learning of the right hand progresses^60,61^. Thus, one possibility is that sequence learning entails a decrease of GABAergic-mediated SICI in the contralateral M1, presumably to facilitate the induction of learning-induced plastic changes^62,63^. To ascertain the relevance of the present results, future work could deliver trains of subthreshold TMS pulses^64^ to disrupt these SICI responses as sequence learning progresses. Doing so would ascertain the causal contribution of the present bilateral SICI responses to motor sequence learning.

Another interesting finding is that these bilateral SICI responses were observed selectively at the onset of motor preparation (i.e., GoCue; Figure 6B-D, also shown in summary Figure 8B), suggesting that sequence learning predominantly altered baseline SICI levels, rather than the bilateral SICI decreases observed during motor preparation *per se*. Although this interpretation aligns with previous ppTMS work showing that motor learning alters resting SICI levels^10,11^, SICI also decreased in the ipsilateral M1 whilst SICI tended to increase in the contralateral M1 (*p* = 0.125) when measured late during preparation (i.e., Late RT; not shown in Figure 8B). This suggests that sequence learning could also alter the bilateral SICI decreases observed during preparation, which merits further investigation. Finally, when considering all the findings in this work, one novel implication is that dynamic changes in bilateral SICI levels contribute to at least two motor processes; an acute decrease of inhibition during motor preparation, and a cooperative but bidirectional shift of baseline inhibition levels as sequence learning progresses. Future work should seek to confirm this.

## Limitations

Here, bilateral CSE and SICI responses were evaluated in separate groups of participants, making it unclear if excitability changes in the ipsilateral M1 correlate to those of the contralateral M1. Future work could assess this by simultaneously evaluating bilateral CSE and SICI using a double-coil TMS paradigm^65,66^. Furthermore, sequence learning extends outside of M1^67,68^, making it unclear how SICI would be altered in other brain areas as sequence learning progresses. Future work could evaluate M1-M1^50^ and/or dorsal premotor cortex (PMd)-M1^69^ corticocortical connectivity using dual-coil TMS^70^ as well as combining TMS with electroencephalography^71^ or magnetic resonance imaging^72^ to address this.

## Conclusion

The present results suggest that bilateral M1s concomitantly decrease SICI during unilateral motor preparation, and that they cooperate to bidirectionally alter baseline SICI levels during motor sequence learning. One possibility is that SICI levels acutely decrease to support motor preparation of various tasks^6–8^, and tonically shift to support motor sequence learning.

## Supporting information

Supplementary Results

## Acknowledgements

This work was funded by the *Fonds de Recherche du Québec – Nature et Technologie* (Québec, Canada), European Research Council Starting Grant (MotMotLearn 637488) and Proof of Concept Grant (ImpHandRehab 872082).

## REFERENCES

1. Robertson, E. M. The serial reaction time task: implicit motor skill learning? J. Neurosci. Off. J. Soc. Neurosci. 27, 10073– 10075 (2007).

2. Schwarb, H. & Schumacher, E. H. Generalized lessons about sequence learning from the study of the serial reaction time task. Adv. Cogn. Psychol. 8, 165–178 (2012).

3. Oliveira, C. M., Hayiou-Thomas, M. E. & Henderson, L. M. The reliability of the serial reaction time task: meta-analysis of test–retest correlations. R. Soc. Open Sci. 10, 221542 (2023).

4. Ariani, G. & Diedrichsen, J. Sequence learning is driven by improvements in motor planning. J. Neurophysiol. 121, 2088– 2100 (2019).

5. Verstynen, T. et al. Dynamic Sensorimotor Planning during Long-Term Sequence Learning: The Role of Variability, Response Chunking and Planning Errors. PLOS ONE 7, e47336 (2012).

6. Reynolds, C. & Ashby, P. Inhibition in the human motor cortex is reduced just before a voluntary contraction. Neurology 53, 730–730 (1999).

7. Dupont-Hadwen, J., Bestmann, S. & Stagg, C. J. Motor training modulates intracortical inhibitory dynamics in motor cortex during movement preparation. Brain Stimulat. 12, 300–308 (2019).

8. Hamel, R. et al. The intracortical excitability changes underlying the enhancing effects of rewards and punishments on motor performance. Brain Stimulat. 16, 1462–1475 (2023).

9. Ziemann, U. et al. TMS and drugs revisited 2014. Clin. Neurophysiol. Off. J. Int. Fed. Clin. Neurophysiol. 126, 1847– 1868 (2015).

10. Coxon, J. P., Peat, N. M. & Byblow, W. D. Primary motor cortex disinhibition during motor skill learning. J. Neurophysiol. 112, 156–164 (2014).

11. Rosenkranz, K., Kacar, A. & Rothwell, J. C. Differential modulation of motor cortical plasticity and excitability in early and late phases of human motor learning. J. Neurosci. Off. J. Soc. Neurosci. 27, 12058–12066 (2007).

12. Lacroix, A. et al. Static magnetic stimulation of the primary motor cortex impairs online but not offline motor sequence learning. Sci. Rep. 9, 9886 (2019).

13. Hashemirad, F., Zoghi, M., Fitzgerald, P. B. & Jaberzadeh, S. The effect of anodal transcranial direct current stimulation on motor sequence learning in healthy individuals: A systematic review and meta-analysis. Brain Cogn. 102, 1–12 (2016).

14. Rumpf, J.-J., May, L., Fricke, C., Classen, J. & Hartwigsen, G. Interleaving Motor Sequence Training With High-Frequency Repetitive Transcranial Magnetic Stimulation Facilitates Consolidation. Cereb. Cortex N. Y. NY 30, 1030– 1039 (2020).

15. Kobayashi, M., Théoret, H. & Pascual-Leone, A. Suppression of ipsilateral motor cortex facilitates motor skill learning. Eur. J. Neurosci. 29, 833–836 (2009).

16. Waters, S., Wiestler, T. & Diedrichsen, J. Cooperation Not Competition: Bihemispheric tDCS and fMRI Show Role for Ipsilateral Hemisphere in Motor Learning. J. Neurosci. 37, 7500–7512 (2017).

17. Narayana, S. et al. Concurrent TMS to the primary motor cortex augments slow motor learning. NeuroImage 85, 971– 984 (2014).

18. Hamano, Y. H., Sugawara, S. K., Fukunaga, M. & Sadato, N. The integrative role of the M1 in motor sequence learning. Neurosci. Lett. 760, 136081 (2021).

19. Cross, E. S., Schmitt, P. J. & Grafton, S. T. Neural Substrates of Contextual Interference during Motor Learning Support a Model of Active Preparation. J. Cogn. Neurosci. 19, 1854–1871 (2007).

20. Cohen, N. R., Cross, E. S., Wymbs, N. F. & Grafton, S. T. Transient disruption of M1 during response planning impairs subsequent offline consolidation. Exp. Brain Res. Exp. Hirnforsch. Exp. Cerebrale 196, 303–309 (2009).

21. Bestmann, S. & Duque, J. Transcranial Magnetic Stimulation: Decomposing the Processes Underlying Action Preparation. Neurosci. Rev. J. Bringing Neurobiol. Neurol. Psychiatry 22, 392–405 (2016).

22. Chye, L., Riek, S., de Rugy, A., Carson, R. G. & Carroll, T. J. Unilateral movement preparation causes task-specific modulation of TMS responses in the passive, opposite limb. J. Physiol. 596, 3725–3738 (2018).

23. Bundy, D. T. & Leuthardt, E. C. The Cortical Physiology of Ipsilateral Limb Movements. Trends Neurosci. 42, 825–839 (2019).

24. Rossi, S., Hallett, M., Rossini, P. M. & Pascual-Leone, A. Screening questionnaire before TMS: an update. Clin. Neurophysiol. Off. J. Int. Fed. Clin. Neurophysiol. 122, 1686 (2011).

25. Lebon, F. et al. Influence of Delay Period Duration on Inhibitory Processes for Response Preparation. Cereb. Cortex 26, 2461–2470 (2016).

26. Soto, O., Valls-Solé, J. & Kumru, H. Paired-Pulse Transcranial Magnetic Stimulation During Preparation for Simple and Choice Reaction Time Tasks. J. Neurophysiol. 104, 1392–1400 (2010).

27. Sinclair, C. & Hammond, G. R. Reduced intracortical inhibition during the foreperiod of a warned reaction time task. Exp. Brain Res. 186, 385–392 (2008).

28. Thomas, C. K., Ross, B. H. & Stein, R. B. Motor-unit recruitment in human first dorsal interosseous muscle for static contractions in three different directions. J. Neurophysiol. 55, 1017–1029 (1986).

29. Caulfield, K. A. et al. Neuronavigation maximizes accuracy and precision in TMS positioning: Evidence from 11,230 distance, angle, and electric field modeling measurements. Brain Stimul. Basic Transl. Clin. Res. Neuromodulation 15, 1192–1205 (2022).

30. Rossi, S., Hallett, M., Rossini, P. M., Pascual-Leone, A., & Safety of TMS Consensus Group. Safety, ethical considerations, and application guidelines for the use of transcranial magnetic stimulation in clinical practice and research. Clin. Neurophysiol. Off. J. Int. Fed. Clin. Neurophysiol. 120, 2008–2039 (2009).

31. Kossev, A. R., Siggelkow, S., Dengler, R. & Rollnik, J. D. Intracortical inhibition and facilitation in paired-pulse transcranial magnetic stimulation: effect of conditioning stimulus intensity on sizes and latencies of motor evoked potentials. J. Clin. Neurophysiol. Off. Publ. Am. Electroencephalogr. Soc. 20, 54–58 (2003).

32. Lefaucheur, J.-P. Transcranial magnetic stimulation. Handb. Clin. Neurol. 160, 559–580 (2019).

33. Chang, W. H. et al. Optimal number of pulses as outcome measures of neuronavigated transcranial magnetic stimulation. Clin. Neurophysiol. Off. J. Int. Fed. Clin. Neurophysiol. 127, 2892–2897 (2016).

34. Biabani, M., Farrell, M., Zoghi, M., Egan, G. & Jaberzadeh, S. The minimal number of TMS trials required for the reliable assessment of corticospinal excitability, short interval intracortical inhibition, and intracortical facilitation. Neurosci. Lett. 674, 94–100 (2018).

35. Smith, V. et al. High-intensity transcranial magnetic stimulation reveals differential cortical contributions to prepared responses. J. Neurophysiol. 121, 1809–1821 (2019).

36. Boisgontier, M. P. & Cheval, B. The anova to mixed model transition. Neurosci. Biobehav. Rev. 68, 1004–1005 (2016).

37. Koerner, T. K. & Zhang, Y. Application of Linear Mixed-Effects Models in Human Neuroscience Research: A Comparison with Pearson Correlation in Two Auditory Electrophysiology Studies. Brain Sci. 7, (2017).

38. Lo, S. & Andrews, S. To transform or not to transform: using generalized linear mixed models to analyse reaction time data. Front. Psychol. 6, (2015).

39. Harrison, X. A. et al. A brief introduction to mixed effects modelling and multi-model inference in ecology. PeerJ 6, e4794 (2018).

40. Benjamini, Y. & Hochberg, Y. Controlling the False Discovery Rate: A Practical and Powerful Approach to Multiple Testing. J. R. Stat. Soc. Ser. B Methodol. 57, 289–300 (1995).

41. Şahin, M. & Aybek, E. Jamovi: An Easy to Use Statistical Software for the Social Scientists. Int. J. Assess. Tools Educ. 6, 670–692 (2020).

42. Chen, R., Cohen, L. G. & Hallett, M. Role of the ipsilateral motor cortex in voluntary movement. Can. J. Neurol. Sci. J. Can. Sci. Neurol. 24, 284–291 (1997).

43. Jenny, A. B. Commissural projections of the cortical hand motor area in monkeys. J. Comp. Neurol. 188, 137–145 (1979).

44. Rouiller, E. M. et al. Transcallosal connections of the distal forelimb representations of the primary and supplementary motor cortical areas in macaque monkeys. Exp. Brain Res. 102, 227–243 (1994).

45. Urrutia-Piñones, J., Morales-Moraga, C., Sanguinetti-González, N., Escobar, A. P. & Chiu, C. Q. Long-Range GABAergic Projections of Cortical Origin in Brain Function. Front. Syst. Neurosci. 16, 841869 (2022).

46. Verstynen, T. & Ivry, R. B. Network Dynamics Mediating Ipsilateral Motor Cortex Activity during Unimanual Actions. J. Cogn. Neurosci. 23, 2468–2480 (2011).

47. Welniarz, Q. et al. The supplementary motor area modulates interhemispheric interactions during movement preparation. Hum. Brain Mapp. 40, 2125–2142 (2019).

48. Estebanez, L., Hoffmann, D., Voigt, B. C. & Poulet, J. F. A. Parvalbumin-Expressing GABAergic Neurons in Primary Motor Cortex Signal Reaching. Cell Rep. 20, 308–318 (2017).

49. Benjamin, P. R., Staras, K. & Kemenes, G. What Roles Do Tonic Inhibition and Disinhibition Play in the Control of Motor Programs? Front. Behav. Neurosci. 4, 30 (2010).

50. Carson, R. G. Inter-hemispheric inhibition sculpts the output of neural circuits by co-opting the two cerebral hemispheres. J. Physiol. 598, 4781–4802 (2020).

51. Buetefisch, C. M., Revill, K. P., Shuster, L., Hines, B. & Parsons, M. Motor demand-dependent activation of ipsilateral motor cortex. J. Neurophysiol. 112, 999–1009 (2014).

52. Berlot, E., Prichard, G., O’Reilly, J., Ejaz, N. & Diedrichsen, J. Ipsilateral finger representations in the sensorimotor cortex are driven by active movement processes, not passive sensory input. J. Neurophysiol. 121, 418–426 (2019).

53. Diedrichsen, J., Wiestler, T. & Krakauer, J. W. Two Distinct Ipsilateral Cortical Representations for Individuated Finger Movements. Cereb. Cortex 23, 1362–1377 (2013).

54. Gielen, S. C. A. M., Schmidt, R. A. & Van Den Heuvel, P. J. M. On the nature of intersensory facilitation of reaction time. Percept. Psychophys. 34, 161–168 (1983).

55. Nickerson, R. S. Intersensory facilitation of reaction time: Energy summation or preparation enhancement? Psychol. Rev. 80, 489–509 (1973).

56. Munzert, J., Lorey, B. & Zentgraf, K. Cognitive motor processes: The role of motor imagery in the study of motor representations. Brain Res. Rev. 60, 306–326 (2009).

57. Hamel, R., Trempe, M. & Bernier, P.-M. Disruption of M1 activity during performance plateau impairs consolidation of motor memories. J. Neurosci. Off. J. Soc. Neurosci. (2017) doi:10.1523/JNEUROSCI.3916-16.2017.

58. Spampinato, D. & Celnik, P. Temporal dynamics of cerebellar and motor cortex physiological processes during motor skill learning. Sci. Rep. 7, 40715 (2017).

59. Orban de Xivry, J.-J., Criscimagna-Hemminger, S. E. & Shadmehr, R. Contributions of the Motor Cortex to Adaptive Control of Reaching Depend on the Perturbation Schedule. Cereb. Cortex 21, 1475–1484 (2011).

60. Kolasinski, J. et al. The dynamics of cortical GABA in human motor learning. J. Physiol. 597, 271–282 (2019).

61. Floyer-Lea, A., Wylezinska, M., Kincses, T. & Matthews, P. M. Rapid modulation of GABA concentration in human sensorimotor cortex during motor learning. J. Neurophysiol. 95, 1639–1644 (2006).

62. Bachtiar, V. & Stagg, C. J. The role of inhibition in human motor cortical plasticity. Neuroscience 278, 93–104 (2014).

63. Barron, H. C. Neural inhibition for continual learning and memory. Curr. Opin. Neurobiol. 67, 85–94 (2021).

64. Nikolov, P. et al. Impact of the number of conditioning pulses on motor cortex excitability: a transcranial magnetic stimulation study. Exp. Brain Res. 239, 583–589 (2021).

65. Grandjean, J. et al. Towards assessing corticospinal excitability bilaterally: Validation of a double-coil TMS method. J. Neurosci. Methods 293, 162–168 (2018).

66. Vassiliadis, P. et al. Using a Double-Coil TMS Protocol to Assess Preparatory Inhibition Bilaterally. Front. Neurosci. 12, 139 (2018).

67. Berlot, E., Popp, N. J. & Diedrichsen, J. A critical re-evaluation of fMRI signatures of motor sequence learning. eLife 9, e55241 (2020).

68. Janacsek, K. et al. Sequence learning in the human brain: A functional neuroanatomical meta-analysis of serial reaction time studies. NeuroImage 207, 116387 (2020).

69. Rosso, C. et al. Anatomical and functional correlates of cortical motor threshold of the dominant hand. Brain Stimulat. 10, 952–958 (2017).

70. Koch, G. Cortico-cortical connectivity: the road from basic neurophysiological interactions to therapeutic applications. Exp. Brain Res. 238, 1677–1684 (2020).

71. Hernandez-Pavon, J. C. et al. TMS combined with EEG: Recommendations and open issues for data collection and analysis. Brain Stimulat. 16, 567–593 (2023).

72. Oathes, D. J. et al. Combining transcranial magnetic stimulation with functional magnetic resonance imaging for probing and modulating neural circuits relevant to affective disorders. Wiley Interdiscip. Rev. Cogn. Sci. 12, e1553 (2021).

