## Supplementary Results for "Bilateral Intracortical Inhibition during Unilateral Motor Preparation and Sequence Learning"

### Supplementary Methods

**Supplementary Table 1 – Number of trials included in the analyses**

**Left M1 group – First experiment**

| | SICI | CSE | NoTMS |
|---|---|---|---|
| **-500ms** | 23.9 ± 0.1 | 23.8 ± 0.2 | 47.8 ± 0.2 |
| **-250ms** | 23.8 ± 0.2 | 23.9 ± 0.1 | |
| **GoCue** | 23.9 ± 0.2 | 23.9 ± 0.2 | |
| **Early RT** | 22.1 ± 1.6 | 22.6 ± 2.5 | |
| **Late RT** | 21.6 ± 1.4 | 21.4 ± 2.3 | |

**Right M1 group – First experiment**

| | SICI | CSE | NoTMS |
|---|---|---|---|
| **-500ms** | 23.7 ± 0.3 | 23.6 ± 0.4 | 47.5 ± 0.7 |
| **-250ms** | 23.5 ± 0.4 | 23.7 ± 0.4 | |
| **GoCue** | 23.6 ± 0.3 | 23.6 ± 0.4 | |
| **Early RT** | 15.5 ± 3.5 | 15.4 ± 3.8 | |
| **Late RT** | 23.4 ± 1.9 | 25.1 ± 1.7 | |

**Left M1 group – Second experiment**

| | | | SICI | CSE | NoTMS |
|---|---|---|---|---|---|
| **Random** | **Early** | **GoCue** | 23.9 ± 0.2 | 23.9 ± 0.2 | 24.0 ± 0.1 |
| | | **Late RT** | 25.5 ± 1.5 | 24.9 ± 1.7 | |
| | **Late** | **GoCue** | 23.5 ± 0.5 | 23.8 ± 0.3 | 23.9 ± 0.1 |
| | | **Late RT** | 26.7 ± 1.0 | 26.8 ± 1.4 | |
| **Learning** | **Early** | **GoCue** | 24.0 ± 0.8 | 23.5 ± 0.4 | 23.7 ± 0.4 |
| | | **Late RT** | 23.8 ± 2.0 | 25.9 ± 2.0 | |
| | **Late** | **GoCue** | 23.6 ± 0.4 | 22.9 ± 0.3 | 24.0 ± 0.1 |
| | | **Late RT** | 22.7 ± 2.0 | 22.0 ± 2.1 | |

**Right M1 group – Second experiment**

| | | | SICI | CSE | NoTMS |
|---|---|---|---|---|---|
| **Random** | **Early** | **GoCue** | 21.2 ± 1.6 | 23.2 ± 1.2 | 23.3 ± 1.2 |
| | | **Late RT** | 25.0 ± 1.9 | 26.1 ± 1.6 | |
| | **Late** | **GoCue** | 21.9 ± 1.5 | 23.4 ± 1.2 | 23.4 ± 1.2 |
| | | **Late RT** | 27.0 ± 1.6 | 28.2 ± 1.5 | |
| **Learning** | **Early** | **GoCue** | 22.2 ± 1.9 | 23.8 ± 0.2 | 23.8 ± 0.4 |
| | | **Late RT** | 23.8 ± 1.2 | 25.8 ± 1.6 | |
| | **Late** | **GoCue** | 21.5 ± 1.5 | 22.9 ± 0.3 | 24.0 ± 0.1 |
| | | **Late RT** | 24.8 ± 1.5 | 23.5 ± 1.5 | |

All descriptive statistics represent the mean ± 95% CIs. The RT rejection criteria and the TMS data (re)pooling procedure slightly decreased the average of CSE and SICI trials at Early RT for the Right M1 group of the first experiment.

#### Supplementary Results – First Experiment (Motor Preparation)

##### TMS over the left and right M1 similarly altered RTs

For the Left M1 group, the RT data (Supplementary Figure 1A) revealed a Time Points * TMS Pulse Types interaction ($\chi^2 = 238.68$, $p < 0.001$), which further revealed simple effects of TMS Pulse Types at each level of Time Points (all $\chi^2 > 7.48$, all $p < 0.030$), except for Late RT ($\chi^2 = 0.59$, $p = 0.745$). The results then revealed that delivering CSE and SICI pulses during the foreperiod (i.e., -500ms, and -250ms) similarly quickened RTs as compared to NoTMS (all $p < 0.001$). At GoCue, only SICI pulses quickened RTs as compared to NoTMS ($p = 0.033$). At Early RT, delivering CSE and SICI pulses after GoCue both slowed RTs as compared to NoTMS (both $p < 0.003$), and CSE further slowed RTs as compared to SICI ($p = 0.002$). This shows that delivering TMS pulses to the contralateral M1 during the foreperiod and at the onset of preparation quickens RTs, whereas delivering TMS pulses early during preparation slows RTs.

For the Right M1 group, the RT data (Supplementary Figure 1C) also revealed a Time Points * TMS Pulse Types interaction ($\chi^2 = 254.47$, $p < 0.001$), which further revealed simple effects of TMS Pulse Types at every level of Time Points (all $\chi^2 > 8.34$, all $p < 0.039$), except at GoCue ($\chi^2 = 4.74$, $p = 0.093$). The results then revealed that delivering CSE and SICI pulses during the foreperiod (i.e., -500ms and -250ms) quickened RTs as compared to NoTMS (all $p < 0.001$). At Early RT, delivering CSE and SICI pulses slowed RTs as compared to NoTMS (both $p < 0.003$). At Late RT, SICI ($p = 0.043$), but not CSE ($p = 0.278$), pulses quickened RTs as compared to NoTMS. Similar to when the left M1 was stimulated, this shows that delivering TMS pulses to the ipsilateral M1 during the foreperiod quickened RTs, whereas delivering TMS pulses early during preparation slowed RTs. Overall, this suggests that bilateral M1s contribute to the processes of motor preparation, as assessed with RTs.

##### TMS over the left M1 selectively slowed MTs

For the Left M1 group, the MT data (Supplementary Figure 1B) revealed a Time Points * TMS Pulse Types interaction ($\chi^2 = 119.49$, $p < 0.001$), which further revealed a simple effect of TMS Pulse Types at Late RT only ($\chi^2 = 63.56$, $p < 0.001$). The results then revealed that CSE ($p < 0.001$) and SICI pulses ($p < 0.001$) both slowed MTs as compared to NoTMS. CSE pulses further slowed MTs as compared to SICI ($p < 0.001$). Overall, this shows that delivering TMS pulses late during preparation slowed MTs.

For the Right M1 group, the MT data (Supplementary Figure 1D) revealed no Time Points * TMS Pulse Types interaction ($\chi^2 = 9.07$, $p = 0.336$), no effect of Time Points ($\chi^2 = 4.12$, $p = 0.390$) and no effect of TMS Pulse Types ($\chi^2 = 0.05$, $p = 0.976$). Unlike when the left M1 was stimulated, this shows that delivering TMS pulses to the ipsilateral M1 late during preparation did not alter MTs. Overall, this suggests that delivering TMS pulses late during preparation – but selectively over the contralateral M1 – disrupts motor execution processes, as assessed with MTs.

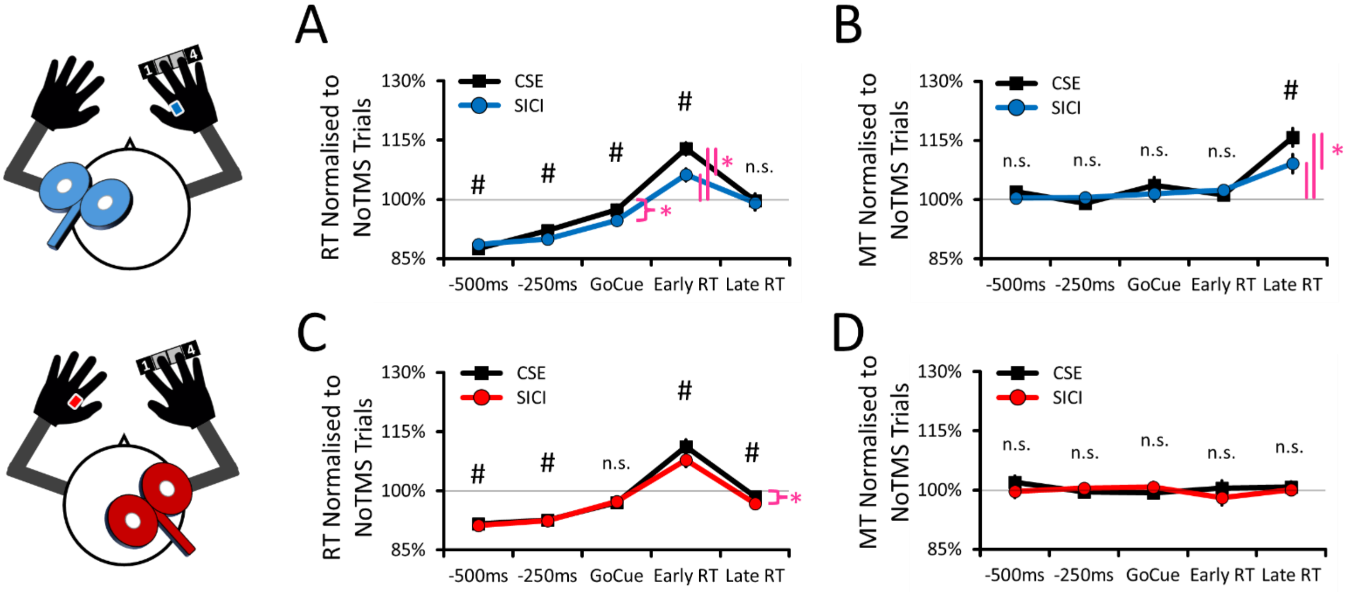


**Supplementary Figure 1. RT and MT data when TMS was delivered to the left and right M1.** For all panels, the RT and MT data have been normalised to the average of NoTMS trials. **(A)** *Left M1 group* – TMS pulses delivered before and at GoCue quickened RT, whereas TMS pulses delivered after GoCue (Early RT only) slowed RTs. **(B)** *Left M1 group* – TMS pulses delivered at Late RT selectively slowed MT. **(C)** *Right M1 group* – TMS pulses to the right M1 altered RT similarly to when the left M1 was stimulated. **(D)** *Right M1 group* – TMS pulses to the right M1 did not disrupt MT. For all panels, the mean ± 95% within-subject CIs are shown. Hashtags (#) indicate significant differences ($p < 0.05$) for both the CSE and SICI trials as compared to NoTMS (100%). The magenta asterisks (*) denote significant differences ($p < 0.05$) for individual conditions as compared to NoTMS (100%).

**TMS over the Left M1 selectively decreased Accuracy**

For the Left M1 group, the Accuracy data (not shown in Supplementary Figure 1) revealed a Time Points * TMS Pulse Types interaction ($\chi^2 = 29.92$, $p < 0.001$), which further revealed simple effects of TMS Pulse Types at GoCue, Early RT, and Late RT (all $p < 0.045$). For all these Time Points, the results then revealed that CSE decreased Accuracy as compared to both SICI (all $p < 0.048$) and NoTMS (all $p < 0.057$). Accuracy on SICI trials did not differ from NoTMS (both $p > 0.489$), except at Late RT where Accuracy decreased on SICI trials as compared to NoTMS ($p < 0.001$). This shows that delivering TMS pulses either at the onset of, as well as early and late during, preparation interfered with accuracy.

For the Right M1 group, the Accuracy data (not shown in Supplementary Figure 2) revealed no Time Points * TMS Pulse Types interaction ($\chi^2 = 1.82$, $p = 0.770$), no effect of Time Points ($\chi^2 = 7.27$, $p = 0.122$), and no effect of TMS Pulse Types ($\chi^2 = 1.24$, $p = 0.265$). Unlike when the left M1 was stimulated, this shows that delivering TMS pulses to the right M1 did not interfere with the ability of participants to accurately execute the sequences with their right hand. Overall, similar to the MT results, this suggests that delivering TMS pulses late during preparation – but selectively over the contralateral M1 – disrupts motor execution processes, as assessed with accuracy.

### Supplementary Results – Second Experiment (Sequence Learning)

#### MTs quickened during sequence learning for both the Left and Right M1 groups

For the Left M1 group, the MT data (Supplementary Figure 2A) revealed an effect of Blocks ($\chi^2 = 14.62$; $p < 0.001$), Phases ($\chi^2 = 5.06$; $p = 0.024$), and TMS Pulse Types ($\chi^2 = 42.84$; $p < 0.001$). See Supplementary Table 4 for a complete report of all results. This shows that MTs were faster on the Learning than on Random blocks ($p < 0.001$). This also shows that MTs quickened from the Early to the Late phase ($p = 0.024$) in both the Learning and Random blocks. The effect of TMS Pulse Types showed that MTs were slower on both CSE ($p < 0.001$) and SICI ($p = 0.005$) trials as compared to NoTMS. MTs on CSE trials were also slower than on SICI ($p < 0.001$). This shows that MTs quickened from the Early to the Late phase in both Random and Learning blocks in the Left M1 group and that delivering TMS pulses over the contralateral M1 slowed MTs.

For the Right M1 group, the MT data (Supplementary Figure 2B) revealed a Blocks * Phases * TMS Pulse Types interaction ($\chi^2 = 7.40$; $p = 0.025$), which was decomposed by conducting simple effects of Phases (Early vs Late) separately for each level of Blocks (Random, Learning) and TMS Pulse Types (NoTMS, CSE, SICI). See Supplementary Table 5 for a complete report of all results. For NoTMS, CSE, and SICI, MTs similarly quickened from Early to Late in the Learning blocks (all $p < 0.035$), but not in the Random ones (all $p > 0.380$). This shows that MTs quickened from the Early to the Late phase only in Learning blocks in the Right M1 group and that delivering TMS pulses over the ipsilateral M1 did not alter MTs of the right hand.

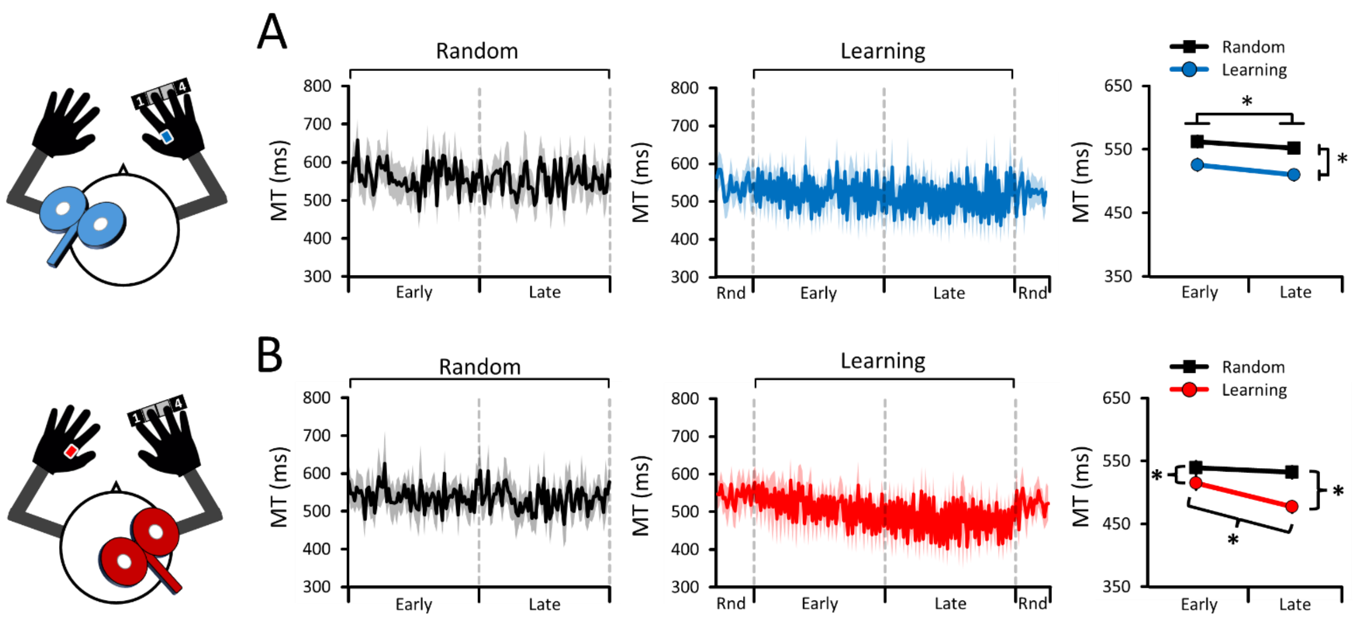


**Supplementary Figure 2. MT data when TMS was delivered to the left and right M1. (A)** *Left M1 group* – MTs similarly quickened in both the Random (pseudorandomised sequences) and Learning blocks (repeating sequences), suggesting that MT non-specifically improved in both blocks. **(B)** *Right M1 group* – MTs quickened in the Learning blocks but not in the Random ones, suggesting that sequence learning improved MTs. For all panels, the mean and 95% within-confidence intervals are shown (either as shaded or whisker error bars). Asterisks (*) denote significant differences ($p < 0.05$). "Rnd" means Random (pseudorandomised sequences).

#### No changes in Accuracy for both the Left and Right M1 groups

For the Left M1 group, the Accuracy data (not shown in Supplementary Figure 2) revealed an effect of TMS Pulse Types ($\chi^2 = 6.01$, $p = 0.050$). See Supplementary Table 6 for a complete report of all results. For TMS Pulse Types, Accuracy decreased on CSE trials as compared to NoTMS ($p = 0.044$). Accuracy on SICI trials did not differ from

CSE ($p$ = 0.256) and NoTMS ($p$ = 0.124). This shows that the learning-specific RT improvements were not accompanied by decreases in accuracy in the Left M1 group and that delivering TMS pulses over the contralateral M1 decreased accuracy for both the Learning and Random blocks.

For the Right M1 group, the Accuracy data (not shown in Supplementary Figure 2) revealed a marginal effect of Blocks ($\chi^2$ = 3.22; $p$ = 0.073). See Supplementary Table 7 for a complete report of all results. For Blocks, Accuracy was marginally greater in Random blocks than in Learning ones ($p$ = 0.073). This shows that the learning-specific RT and MT improvements were not accompanied by decreases in accuracy in the Right M1 group and that TMS pulses over the ipsilateral M1 did not alter the accuracy of the right hand.

| **Supplementary Table 2 – Left M1 group – Second experiment – RT Results** | | |
|---|---|---|
| Blocks | $\chi^2$ = 15.52 | $p$ < 0.001 |
| Phases | $\chi^2$ = 16.18 | $p$ < 0.001 |
| TMS Pulse Types | $\chi^2$ = 14.67 | $p$ = 0.001 |
| **Blocks * Phases** | $\chi^2$ = 12.39 | $p$ < 0.001 |
| Blocks * TMS Pulse Types | $\chi^2$ = 4.23 | $p$ = 0.121 |
| Phases * TMS Pulse Types | $\chi^2$ = 1.06 | $p$ = 0.587 |
| Blocks * Phases * TMS Pulse Types | $\chi^2$ = 0.66 | $p$ = 0.718 |

| **Supplementary Table 3 – Right M1 group – Second experiment – RT Results** | | |
|---|---|---|
| Blocks | $\chi^2$ = 40.43 | $p$ < 0.001 |
| Phases | $\chi^2$ = 30.20 | $p$ < 0.001 |
| TMS Pulse Types | $\chi^2$ = 58.77 | $p$ < 0.001 |
| **Blocks * Phases** | $\chi^2$ = 74.52 | $p$ < 0.001 |
| **Blocks * TMS Pulse Types** | $\chi^2$ = 6.17 | $p$ = 0.046 |
| Phases * TMS Pulse Types | $\chi^2$ = 0.47 | $p$ = 0.792 |
| Blocks * Phases * TMS Pulse Types | $\chi^2$ = 3.68 | $p$ = 0.159 |

| **Supplementary Table 4 – Left M1 group – Second experiment – MT Results** | | |
|---|---|---|
| **Blocks** | $\chi^2$ = 14.62 | $p$ < 0.001 |
| **Phases** | $\chi^2$ = 5.06 | $p$ = 0.024 |
| **TMS Pulse Types** | $\chi^2$ = 42.83 | $p$ < 0.001 |
| Blocks * Phases | $\chi^2$ = 2.16 | $p$ = 0.142 |
| Blocks * TMS Pulse Types | $\chi^2$ = 0.14 | $p$ = 0.934 |
| Phases * TMS Pulse Types | $\chi^2$ = 0.74 | $p$ = 0.690 |
| Blocks * Phases * TMS Pulse Types | $\chi^2$ = 5.11 | $p$ = 0.078 |

| **Supplementary Table 5 – Right M1 group – Second experiment – MT Results** | | |
|---|---|---|
| Blocks | $\chi^2$ = 14.69 | $p$ < 0.001 |
| Phases | $\chi^2$ = 18.93 | $p$ < 0.001 |
| TMS Pulse Types | $\chi^2$ = 1.80 | $p$ = 0.406 |
| Blocks * Phases | $\chi^2$ = 6.88 | $p$ = 0.009 |
| Blocks * TMS Pulse Types | $\chi^2$ = 14.12 | $p$ < 0.001 |
| Phases * TMS Pulse Types | $\chi^2$ = 11.51 | $p$ = 0.003 |
| Blocks * Phases * TMS Pulse Types | $\chi^2$ = 7.40 | $p$ = 0.025 |

| Supplementary Table 6 – Left M1 group – Second experiment – Accuracy Results | | |
|---|---|---|
| Blocks | $\chi^2 = 1.60$ | $p = 0.206$ |
| Phases | $\chi^2 = 1.65$ | $p = 0.198$ |
| **TMS Pulse Types** | $\chi^2 = 6.01$ | $p = 0.050$ |
| Blocks * Phases | $\chi^2 < 0.01$ | $p = 0.976$ |
| Blocks * TMS Pulse Types | $\chi^2 = 1.10$ | $p = 0.577$ |
| Phases * TMS Pulse Types | $\chi^2 = 2.25$ | $p = 0.325$ |
| Blocks * Phases * TMS Pulse Types | $\chi^2 = 1.98$ | $p = 0.372$ |

| Supplementary Table 7 – Right M1 group – Second experiment – Accuracy Results | | |
|---|---|---|
| Blocks | $\chi^2 = 3.22$ | $p = 0.073$ |
| Phases | $\chi^2 = 0.26$ | $p = 0.608$ |
| TMS Pulse Types | $\chi^2 = 4.70$ | $p = 0.096$ |
| Blocks * Phases | $\chi^2 = 2.02$ | $p = 0.156$ |
| Blocks * TMS Pulse Types | $\chi^2 = 2.42$ | $p = 0.298$ |
| Phases * TMS Pulse Types | $\chi^2 = 3.34$ | $p = 0.189$ |
| Blocks * Phases * TMS Pulse Types | $\chi^2 = 1.09$ | $p < 0.580$ |

| Supplementary Table 8 – Left M1 group – Second experiment – CSE Results | | |
|---|---|---|
| Blocks | $\chi^2 = 28.76$ | $p < 0.001$ |
| Phases | $\chi^2 = 0.75$ | $p = 0.387$ |
| Time Points | $\chi^2 = 5.51$ | $p = 0.019$ |
| Initiating Fingers | $\chi^2 = 87.28$ | $p < 0.001$ |
| Blocks * Phases | $\chi^2 = 5.78$ | $p = 0.016$ |
| Blocks * Time Points | $\chi^2 = 7.87$ | $p = 0.005$ |
| Phases * Time Points | $\chi^2 = 0.51$ | $p = 0.474$ |
| Blocks * Initiating Fingers | $\chi^2 = 0.88$ | $p = 0.347$ |
| Phases * Initiating Fingers | $\chi^2 = 1.16$ | $p = 0.282$ |
| Time Points * Initiating Fingers | $\chi^2 = 86.35$ | $p < 0.001$ |
| Blocks * Phases * Time Points | $\chi^2 = 1.92$ | $p = 0.166$ |
| Blocks * Phases * Initiating Fingers | $\chi^2 = 1.20$ | $p = 0.272$ |
| **Blocks * Time Points * Initiating Fingers** | $\chi^2 = 26.08$ | $p < 0.001$ |
| Phases * Time Points * Initiating Fingers | $\chi^2 < 0.01$ | $p = 0.984$ |
| Blocks * Phases * Time Points * Initiating Fingers | $\chi^2 = 1.29$ | $p = 0.256$ |

| Supplementary Table 9 – Right M1 group – Second experiment – CSE Results | | |
|---|---|---|
| Blocks | $\chi^2 = 0.69$ | $p = 0.407$ |
| Phases | $\chi^2 = 0.98$ | $p = 0.322$ |
| Time Points | $\chi^2 = 7.01$ | $p = 0.008$ |
| Initiating Fingers | $\chi^2 = 11.23$ | $p = 0.001$ |
| Blocks * Phases | $\chi^2 = 4.00$ | $p = 0.046$ |
| Blocks * Time Points | $\chi^2 = 7.41$ | $p = 0.007$ |
| Phases * Time Points | $\chi^2 = 0.96$ | $p = 0.326$ |
| Blocks * Initiating Fingers | $\chi^2 = 1.26$ | $p = 0.255$ |
| Phases * Initiating Fingers | $\chi^2 = 4.50$ | $p = 0.034$ |
| Time Points * Initiating Fingers | $\chi^2 = 22.20$ | $p < 0.001$ |
| Blocks * Phases * Time Points | $\chi^2 = 2.65$ | $p = 0.103$ |
| Blocks * Phases * Initiating Fingers | $\chi^2 = 0.19$ | $p = 0.664$ |
| **Blocks * Time Points * Initiating Fingers** | $\chi^2 = 3.82$ | $p = 0.051$ |
| Phases * Time Points * Initiating Fingers | $\chi^2 = 1.58$ | $p = 0.208$ |
| Blocks * Phases * Time Points * Initiating Fingers | $\chi^2 = 0.07$ | $p = 0.793$ |

| Supplementary Table 10 – Left M1 group – Second experiment – SICI Results | | |
|---|---|---|
| Blocks | $\chi^2 = 1.67$ | $p = 0.196$ |
| Phases | $\chi^2 = 0.95$ | $p = 0.330$ |
| Time Points | $\chi^2 = 9.74$ | $p = 0.002$ |
| Initiating Fingers | $\chi^2 = 54.63$ | $p < 0.001$ |
| Blocks * Phases | $\chi^2 = 0.05$ | $p = 0.819$ |
| Blocks * Time Points | $\chi^2 < 0.01$ | $p = 0.959$ |
| Phases * Time Points | $\chi^2 = 5.47$ | $p = 0.019$ |
| Blocks * Initiating Fingers | $\chi^2 = 0.10$ | $p = 0.751$ |
| Phases * Initiating Fingers | $\chi^2 = 0.39$ | $p = 0.530$ |
| Time Points * Initiating Fingers | $\chi^2 = 63.75$ | $p < 0.001$ |
| **Blocks * Phases * Time Points** | $\chi^2 = 3.94$ | $p = 0.047$ |
| Blocks * Phases * Initiating Fingers | $\chi^2 = 0.61$ | $p = 0.436$ |
| **Blocks * Time Points * Initiating Fingers** | $\chi^2 = 6.37$ | $p = 0.012$ |
| Phases * Time Points * Initiating Fingers | $\chi^2 = 0.30$ | $p = 0.582$ |
| Blocks * Phases * Time Points * Initiating Fingers | $\chi^2 = 0.38$ | $p = 0.539$ |

| Supplementary Table 11 – Right M1 group – Second experiment – SICI Results | | |
|---|---|---|
| Blocks | $\chi^2 = 0.82$ | $p = 0.366$ |
| Phases | $\chi^2 = 1.25$ | $p = 0.264$ |
| Time Points | $\chi^2 = 8.07$ | $p = 0.004$ |
| Initiating Fingers | $\chi^2 = 20.18$ | $p < 0.001$ |
| Blocks * Phases | $\chi^2 = 0.58$ | $p = 0.447$ |
| Blocks * Time Points | $\chi^2 = 0.06$ | $p = 0.808$ |
| Phases * Time Points | $\chi^2 = 2.41$ | $p = 0.121$ |
| Blocks * Initiating Fingers | $\chi^2 = 0.76$ | $p = 0.384$ |
| Phases * Initiating Fingers | $\chi^2 = 2.72$ | $p = 0.099$ |
| **Time Points * Initiating Fingers** | $\chi^2 = 16.29$ | $p < 0.0001$ |
| **Blocks * Phases * Time Points** | $\chi^2 = 8.89$ | $p = 0.0029$ |
| Blocks * Phases * Initiating Fingers | $\chi^2 = 0.63$ | $p = 0.426$ |
| Blocks * Time Points * Initiating Fingers | $\chi^2 < 0.01$ | $p = 0.990$ |
| Phases * Time Points * Initiating Fingers | $\chi^2 = 0.06$ | $p = 0.800$ |
| Blocks * Phases * Time Points * Initiating Fingers | $\chi^2 = 1.53$ | $p = 0.216$ |
